# Dnmt3a-dependent *de novo* DNA methylation enforces lineage commitment and preserves functionality of memory Th1 and Tfh cells

**DOI:** 10.1101/2024.12.03.623450

**Authors:** Bryant Perkins, Camille L. Novis, Andrew Baessler, Linda M. Sircy, Monyca M. Thomas, Malia Harrison-Chau, Andrew W. Richens, Bryce Fuchs, Nguyen X. Nguyen, Kaitlyn Flint, Brittany M. Strobelt, Katherine E. Varley, J. Scott Hale

**Affiliations:** Division of Microbiology and Immunology, Department of Pathology, University of Utah School of Medicine, Salt Lake City, UT 84112; Department of Oncological Sciences, Huntsman Cancer Institute, University of Utah School of Medicine, Salt Lake City, UT 84112, USA

## Abstract

Following acute viral infection, naïve CD4+ T cells differentiate into T follicular helper (Tfh) and T helper 1 (Th1) cells that generate long-lived memory cells. However, it is unclear how memory Tfh and Th1 cells maintain their lineage commitment. We demonstrate that Tfh and Th1 lineages acquire distinct Dnmt3a-dependent *de novo* DNA methylation programs that are preserved into memory. *Dnmt3a* deletion impairs lineage commitment and functionality of memory Th1 and Tfh cells, resulting in aberrant *Runx1* upregulation that represses germinal center Tfh cell differentiation. In contrast, transient pharmacological DNA methyltransferase inhibition during priming impairs repression of Tfh-associated genes while properly silencing *Runx1*, and results in enhanced Tfh cell functionality in primary and secondary responses to viral infections. Together, these findings demonstrate that Dnmt3a-mediated epigenetic programing is required to enforce T helper lineage commitment and preserve Tfh and Th1-specific functions during the recall response to infection, and reveal novel strategies to improve long-lived adaptive immunity against infectious diseases.

**SUMMARY:** This article demonstrates that Dnmt3a-dependent epigenetic programing regulates functionality and plasticity of Th1 and Tfh memory cells. Furthermore, early pharmacological inhibition of such programing enhances GC Tfh cell differentiation, suggesting novel strategies for modulating the immune response to viral infections.

## INTRODUCTION

In response to acute viral infections, the immune system utilizes humoral and cellular mediated mechanisms to clear pathogens and provide lasting protection against reinfection. CD4+ T helper cells are critical in both aspects. T helper 1 (Th1) cells migrate to sites of infection where they coordinate cell-mediated immune responses (Zhu et al., 2010). T follicular helper (Tfh) cells are critical regulators of the humoral immune response that are required for formation of the germinal center (GC), a transient substructure of the B cell follicle where GC B cells undergo clonal selection, affinity maturation, and differentiation into memory B cells and plasma cells (Crotty, 2019). Following antigen clearance, the majority of effector T helper cells undergo apoptosis, leaving behind a population of T helper cells that persist to form long-lived memory (Künzli and Masopust, 2023). Both Tfh and Th1 cells have the capacity to form long-lived memory cells that preferentially recall their specialized effector functions upon viral reinfection (MacLeod et al., 2011, Marshall et al., 2011, Morita et al., 2011, Pepper et al., 2011, Lüthje et al., 2012, Liu et al., 2012, Weber et al., 2012, Hale et al., 2013, Choi et al., 2013, Künzli et al., 2020, Baessler et al., 2023, Feng et al., 2024).

Increasing humoral immune responses may improve vaccine efficacy against viral infections, including influenza virus and HIV (Burton et al., 2012, Krammer and Palese, 2015). Several studies have demonstrated that the quantity of GC Tfh cell and memory Tfh cells positively correlates with the magnitude of the germinal response and production of neutralizing antibodies to viral infections (Locci et al., 2013, Havenar-Daughton et al., 2016, Cirelli et al., 2019, Lee et al., 2021). Thus, developing strategies to enhance the abundance and quality of Tfh cells may improve long-lasting antibody-mediated protection against infectious disease.

Tfh cells are defined by expression of transcription factor Bcl6 and the chemokine receptor CXCR5 (Johnston et al., 2009, Nurieva et al., 2009, Yu et al., 2009). Early Tfh differentiation is driven by the factors Tcf1 and Lef1, which directly promotes Bcl6 expression (Wu et al., 2015, Choi et al., 2015, Xu et al., 2015, Shao et al., 2019). Several other transcription factors repress Tfh differentiation, including Blimp1, Foxo1, Runx2, and Runx3 (Johnston et al., 2009, Stone et al., 2015, Choi and Crotty, 2021, Baessler et al., 2022, Lai et al., 2022). In addition to transcription factors that repress or activate gene expression, epigenetic reprograming of the cell drives differentiation and establishes heritable transcriptional programing. This includes DNA methylation, an epigenetic program whereby methylation is added to CpG dinucleotides at regulatory regions to silence gene expression (Greenberg and Bourc’his, 2019). A growing body of research has demonstrated that T helper differentiation is accompanied by changes in DNA methylation (Yu et al., 2012, Gamper et al., 2009, Thomas et al., 2012, Hale et al., 2013, Pham et al., 2013, Tsagaratou et al., 2014, Ichiyama et al., 2015, Yue et al., 2016, Yue et al., 2019, Yue et al., 2021, Baessler et al., 2022, Baessler et al., 2023). For example, active DNA demethylation mediated by the methylcytosine dioxygenase Tet2, inhibits GC Tfh cell and memory Tfh cell formation during viral infection in coordination with the transcription factors Foxo1 and *Runx1* (Baessler et al., 2022, Baessler et al., 2023). The DNA methyltransferase Dnmt3a catalyzes *de novo* methylation during T helper differentiation and has been implicated in regulating T cell function by restraining cytokine production in Th1 and Th2 cells and in silencing genes associated with alternative T helper lineages (Gamper et al., 2009, Thomas et al., 2012, Yu et al., 2012). However, whether Dnmt3a and *de novo* methylation regulates Tfh and Th1 cell differentiation, function, memory formation, and lineage commitment is poorly understood.

In this study, we examined the role of Dnmt3a-dependent *de novo* methylation programing in Tfh and Th1 cell differentiation, function, and lineage commitment. We report that Tfh and Th1 cells acquire distinct Dnmt3a-mediated lineage-specific *de novo* methylation programs that are maintained in memory Tfh and Th1 cells. Using the methyltransferase inhibitor Decitabine, we demonstrate that transient inhibition of Dnmt3a activity early during T cell priming reduces memory Th1 cell lineage commitment while also improving Tfh cell functionality to primary and secondary viral infections, representing a novel approach to enhance germinal center responses. Deletion of *Dnmt3a* similarly resulted in memory Th1 cells that were less lineage committed; however, *Dnmt3a* deficient secondary Tfh cells were functionally impaired, being unable to form germinal center Tfh cells during the memory recall response. We attribute this to loss of *de novo* methylation and aberrant upregulation of *Runx1* in Dnmt3a-deficient secondary Tfh cells, and further, we demonstrate that *Runx1* overexpression represses GC Tfh differentiation. Importantly, Decitabine treatment during T cell priming did not inhibit silencing of *Runx1,* but did partially inhibit Dnmt3a-mediated programing at several Tfh-associated genes in Tfh and Th1 cells. Thus, early inhibition of Dnmt3a-mediated *de novo* methylation using decitabine can enhance the formation of highly functional Tfh cell and memory Tfh cells and may be a strategy to improve the germinal center response to viral infections. Together, our data demonstrates a critical role of Dnmt3a-mediated *de novo* methylation for promoting T helper lineage commitment and functionality in memory CD4+ T cells and also indicates that partial inhibition of DNA methylation programing could improve the GC Tfh cell response to viral pathogens.

## RESULTS

### Lineage-specific *de novo* methylation programing is acquired during primary differentiation and maintained in memory Tfh and Th1 cells

To address whether *de novo* methylation programing underpins effector and memory Tfh and Th1 differentiation, we used the acute lymphocytic choriomeningitis virus (LCMV) infection model. Naive SMARTA TCR transgenic CD4+ T cells (CD45.1+) that are specific for the LCMV GP^66-77^ epitope were adoptively transferred into C57BL/6J (B6) mice (Oxenius et al., 1998). Recipient mice were subsequently infected with LCMV. At 7 and 60+ days post infection, Tfh (CXCR5+ Ly6C^LO^) and Th1 (CXCR5-Ly6C^HI^) cells were sorted to isolate genomic DNA for whole genome DNA methylation analysis. Differentially methylated regions (DMRs) were identified for effector or memory Tfh and Th1 cells relative to naïve T cells.

This analysis revealed genome wide changes in methylation programing at over 25k regions per cell type relative to naïve CD4+ T cells (Fig. 1A). *De novo* methylation accounted for a small fraction of these DMRs (Fig. 1A).

**Figure 1.**
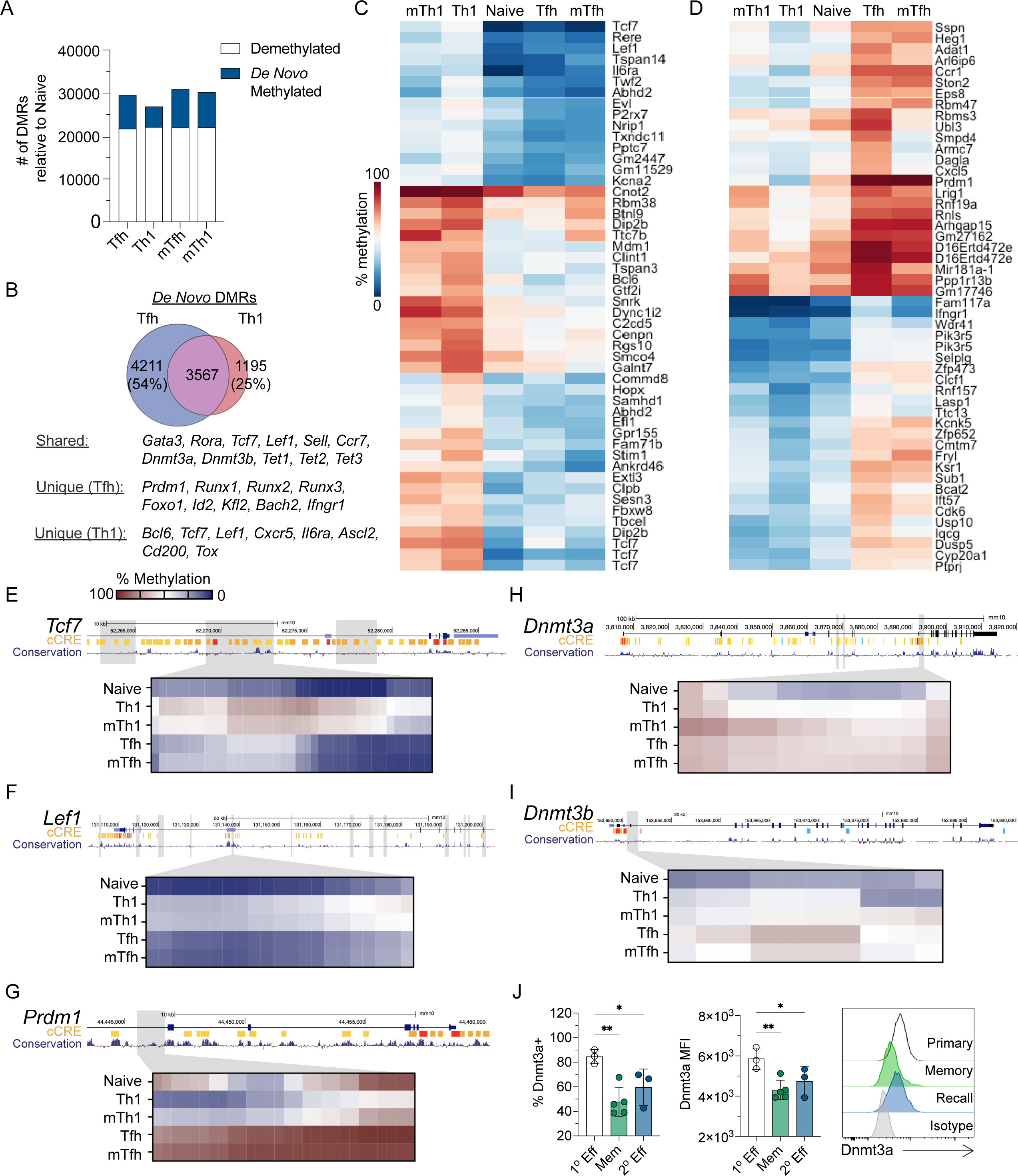
Lineage-specific *de novo* methylation programing is acquired during primary differentiation and maintained in memory Tfh and Th1 cells. Whole genome bisulfite sequencing was performed on DNA isolated from Tfh and Th1 cells sorted at 7(effector) or 60+ days (memory) post LCMV infection. DMRs were identified between each cell type relative to naïve CD4+ T cells. Data are representative of two biological replicates. (**A**) Quantification of the differentially methylated regions (DMRs) by cell type, broken down as *de novo* methylated (hypermethylated) or demethylated (hypomethylated) relative to naïve CD4+ T cells. (**B**) Venn diagram of overlapping (shared) *de novo* DMRs between effector Tfh and Th1 cells. (C-D) Heatmaps of the top 50 lineage-specific *de novo* DMRs for (**C**) Th1 or (**D**) Tfh cells. DMRs were ranked by mean difference in methylation between Tfh and Th1 cells. (E-I) UCSC genome browser snapshots of (**E**) *Tcf7*, (**F**) *Lef1*, (**G**) *Prdm1*, (**H**) *Dnmt3a*, and (**I**) *Dnmt3b.* Tracks show ENCODE identified candidate cis regulatory elements (cCREs) and mammalian conservation. Grey boxes identify *de novo* DMRs. Select DMRs are highlighted in the heatmaps below and each column represents a single CpG site within the region. Significant DMRs were identified using BSmooth DMR finder and have at least 10 CpGs and an absolute mean difference of >0.1. (J) Flow cytometric analysis was performed on primary effector SMARTA cells (Day 7), memory SMARTA cells (Day 42), or secondary effector SMARTA cells (Secondary Day 7) following primary or secondary LCMV infections. For secondary effector analysis, SMARTA cells were isolated from the spleens of LCMV memory mice, CD4+ T cells were magnetically enriched, and cells were transferred into naïve recipients. All data is from analysis of the spleen. Data are representative of two independent experiments. (**J**) Frequency of Dnmt3a+ SMARTA cells and Dnmt3a MFI of SMARTA cells are shown as are representative Dnmt3a histograms. Statistical significance was determined by an unpaired t test (* p<0.05, **p<0.01).

During LCMV infection, Tfh cells partially take on Th1 cell transcriptional qualities, including intermediate expression of Tbet (Hale et al., 2013). Therefore, we reasoned that aspects of *de novo* methylation programing may be shared and that the unique programing would be foundational to Tfh and Th1 cell identity and lineage commitment. To determine shared *de novo* methylation programs of Tfh and Th1 effector cells, we first identified overlapping DMRs. Tfh cells shared 46% of *de novo* DMRs with Th1 cells, while 75% of Th1 *de novo* DMRs were shared with Tfh cells (Fig. 1B). Shared *de novo* DMRs were found at genes associated with Th2/Th17 cells (*Gata3, Rora*), memory formation (*Sell, Ccr7, Bcl2*), and DNA methylation (*Dnmt3a, Dnmt3b, Tet2*). We next ranked *de novo* DMRs based on mean difference in methylation between Tfh and Th1 cells. Our heatmap analysis of the top 50 Th1-specific DMRs highlighted several Tfh genes which acquired *de novo* methylation (*Tcf7, Lef1, Bcl6*, *P2rx7,* and *Il6ra*) (Fig. 1C). The top 50 Tfh-specific *de novo* DMRs included the Th1 genes *Prdm1, Ifngr1*, and *Selplg* (Fig. 1D). We further analyzed the genomic locations of select Th1 DMRs and found evidence of extensive intragenic *de novo* methylation at *Tcf7* and *Lef1* that overlapped with regions of mammalian conservation (Fig. 1E-F). In Tfh cells, we found clear evidence of *de novo* methylation within the *Prdm1* locus (Fig. 1G). Of note, this Tfh *de novo* DMR underwent demethylation in the Th1 lineage which express Blimp1. Overall, these analyses demonstrates that effector and memory Tfh and Th1 cells acquire unique *de novo* methylation programing which may be linked to lineage commitment.

These analyses also revealed that the *de novo* methylation programs were shared between effector and memory T cells of the same lineage (Fig. 1C-I), indicating that lineage-specific *de novo* methylation programing was acquired during T cell differentiation and then maintained into memory. In agreement with this idea, we observed *de novo* DMRs at the genes *Dnmt3a* and *Dnmt3b* (Fig. 1H-I). To determine whether *de novo* methylation at the *Dnmt3a* locus correlated with reduced expression in memory and secondary (2°) effector T cells, we measured Dnmt3a by flow cytometry. For primary effector or memory timepoints, SMARTA T cells were adoptively transferred into B6 mice and infected with LCMV. For memory recall experiments, SMARTA T cells were enriched at memory timepoint and transferred into naïve B6 recipients, which were then infected with LCMV. At the primary effector timepoint, we found that greater than 80% of SMARTA T cells expressed Dnmt3a (Fig. 1J). In contrast, there was a significant reduction in the percent of SMARTA cells that expressed Dnmt3a at the memory and recall timepoints (Fig. 1J). These data suggest that *de novo* methylation programing is largely acquired during the primary infection.

### Early DNA methyltransferase inhibition using decitabine during T cell priming enhances germinal center Tfh differentiation to primary viral infections

Given the unique *de novo* methylation programs that are acquired during Tfh and Th1 cell differentiation, we reasoned that inhibition of *de novo* methylation programing would alter T helper differentiation and function. To address this, we first used an inhibitor of DNA methyltransferases, Decitabine (5’-Aza-2’-deoxycitidine, DAC) (Derissen et al., 2013, Seelan et al., 2018, Christman, 2002). Dnmt3a is upregulated within 24 hours *in vitro* (Gamper et al., 2009, Thomas et al., 2012), so we reasoned that DAC treatment would need to occur early to inhibit *de novo* methylation. DAC is also cytotoxic at high concentrations (Christman, 2002, Derissen et al., 2013, Seelan et al., 2018). Therefore, we first performed *in vivo* titrations of DAC to find a dose that would not impair T and B cell responses. At doses of 0.75 mg/kg, we found no difference in the number of B cells or T cells relative to PBS control (Fig S1A-B), suggesting this dose did not induce cell death. Intriguingly, we did find evidence of increased GC Tfh cell differentiation, even at higher cytotoxic doses (Fig. S1C). In follow up experiments, naïve SMARTA cells were transferred into B6 recipient mice, infected with LCMV, and treated at 20 hours post infection with either DAC (0.75mg/kg) or PBS (Fig. 2A). DAC did not alter SMARTA T effector cell accumulation or Tfh/Th1 differentiation (Fig. 2B-C). However, DAC treatment significantly increased the percent and number of Bcl6^HI^ GC Tfh cells (Fig. 2D-E) and Bcl6 was upregulated in DAC treated Tfh/Th1 cells (Fig. 2F). In a follow up experiment, we analyzed the effect of DAC treatment on the polyclonal T cell response to LCMV. This analysis demonstrated that DAC treatment enhanced polyclonal PD-1^HI^ GC Tfh cell responses and resulted in more GC B cells (Fig. S1D-E). Thus, early decitabine treatment during T cell priming enhanced Bcl6 expression and GC Tfh cell differentiation during LCMV infection.

**Figure 2.**
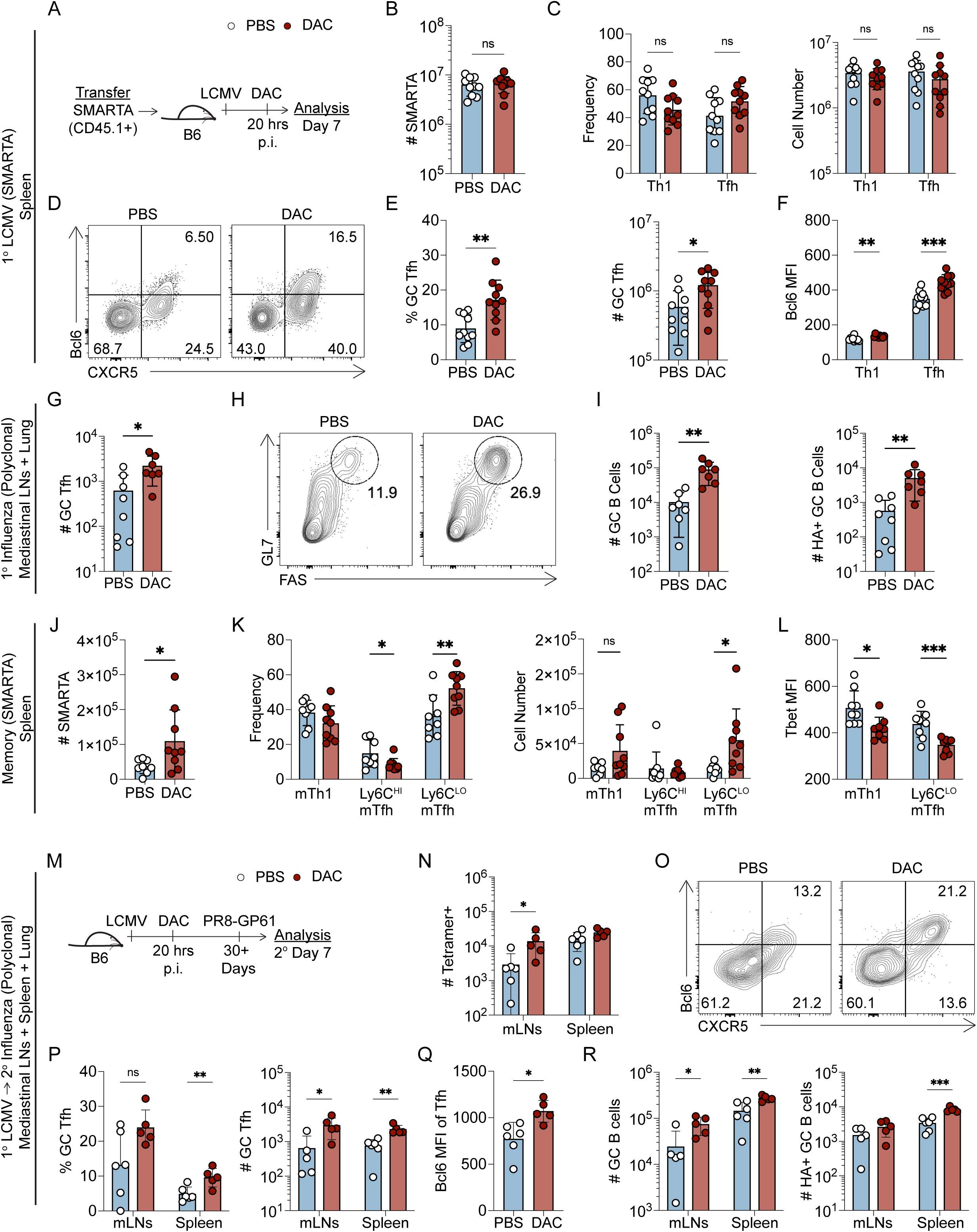
Early decitabine treatment during T cell priming enhances germinal center Tfh differentiation to primary and secondary viral infections. (A-F) Naïve SMARTA cells were transferred into B6 mice which were subsequently infected with LCMV. At 20 hours post infection, mice received an injection of either DAC (0.75 mg/kg) or PBS. Analysis was performed on the spleen at 7 days post infection. Data are representative of more than three experiments. (**A**) experimental schematic. (**B**) Total number of SMARTA cell. (**C**) Percent and number of CXCR5+ Tfh cells or CXCR5-Tfh cells, gated on SMARTA cells. (**D**) Representative FACS plot of CXCR5 and Bcl6 for identification of GC Tfh, gated on SMARTA T cells. (**E**) Percent and number of CXCR5+ Bcl6^HI^ GC Tfh SMARTA cells. (**F**) Bcl6 MFI of Tfh and Th1 SMARTA cells. Statistical significance was determined by an unpaired t Test (* p<0.05, **p<0.01, ***p<0.001). (G-I) B6 mice were infected with the influenza strain PR8 and given either a single dose of DAC (0.375 mg/kg) or PBS at 20 hours post infection. Analysis was performed on the mediastinal lymph nodes (mLNs) at 8 days post infection. Data are representative of two experiments. (**G**) Quantification of the total numbers of CXCR5+ Bcl6^hi^ GC Tfh cells, gated on NP-specific CD4+ T cells. (**H**) Representative FACS plot GL7 and FAS for identification of GC B cells, gated on IgD negative B cells. (**I**) Total number of GC B cells and hemagglutinin-specific GC B cells. Statistical significance was determined by an unpaired t Test (* p<0.05, **p<0.01, ***p<0.001). (J-L) Naïve SMARTA cells were transferred into B6 mice which were subsequently infected with LCMV. At 20 hours post infection, mice received an injection of either DAC (0.75 mg/kg) or PBS. Analysis was performed on the spleen approximately 60 days later. Data are representative of more than three experiments. (**J**) Total number of SMARTA cells. (**K**) Percent and number of CXCR5-Ly6C^hi^ mTh1, CXCR5+ Ly6C^hi^ mTfh, and CXCR5+ Ly6C^lo^ mTfh cells. (**L**) The MFI of Tbet was quantified for CXCR5-Ly6C^hi^ mTh1 and CXCR5+ Ly6C^lo^ mTfh cells. Statistical significance was determined by an unpaired t Test (* p<0.05, **p<0.01, ***p<0.001). (M-R) B6 mice were infected with LCMV and given a single dose of DAC (0.75mg/kg) 20 hours p.i. At 30+ days p.i., mice were intranasally infected with 500 TCID_50_ of recombinant influenza PR8-GP^61-80^. The following issues were harvested for analysis at 7 days post influenza infection: mediastinal LNs (mLNs), spleen (sp), and lung. Data are representative of two experiments. (**M**) Experimental setup. (**N**) Total number of GP-specific CD4+ T cells. (**O**) Representative FACS plots of Bcl6 and CXCR5 for GC Tfh identification, gated on GP-specific CD4 T cells. (**P**) Percent and number of CXCR5+ Bcl6^hi^ GC Tfh cells. (**Q**) The MFI of Bcl6 was quantified from CXCR5+ Tfh cells. (**R**) Number of GC B cells and hemagglutinin-specific GC B cells. Statistical significance was determined by an unpaired t Test (* p<0.05, **p<0.01, ***p<0.001).

To determine if DAC treatment enhanced the GC Tfh cell responses in other viral infection models, mice were intranasally infected with 200 TCID_50_ of the Influenza strain PR8 and treated with DAC (0.375 mg/kg) at 20 hours post infection. For the influenza model, we used this dose of DAC because we observed signs of cytotoxicity at the 0.75 mg/kg. We analyzed the T cell response 8 days later and found a significant increase in NP-specific CD4+ T cells in the mediastinal lymph nodes (mLNs) but not the lung of DAC treated recipients (Fig. S1F). In the influenza model, DAC treatment enhanced both Tfh and Bcl6^HI^ GC Tfh differentiation in the mLNs (Fig. 2G, S1G). We also observed a significant increase in the total number of GC B cells as well as hemagglutinin-specific GC B cells (Fig. 2H-I).

Importantly, T regulatory responses were unaffected by DAC treatment in the mLNs (Fig. S1H). Thus, these data indicate that early DAC treatment enhances GC Tfh differentiation and function to multiple viral infections.

### Decitabine treatment during T cell priming enhances memory Tfh cell formation

We examined the effects of early (day 1 during primary response) DAC treatment on memory cell formation by analyzing the spleen at day 60 post infection. Early DAC treatment enhanced the numbers of total SMARTA memory cells as well as the Ly6C^LO^ subset of memory Tfh (mTfh) cells (Fig. 2J-K). Importantly, while there was no effect on memory Th1 cell (mTh1) accumulation (Fig. 2K), both mTh1 and mTfh cells expressed less Tbet in DAC-treated recipients (Fig. 2L). These data demonstrate that early DAC treatment during the primary response enhances memory Tfh cell formation in the spleen.

### Decitabine treatment during T cell priming enhances functional memory CD4+ T cell recall responses to heterologous infection with Influenza

Based on the previous results, we hypothesized that DAC treatment will result in memory cells that are intrinsically programmed to preferentially form GC Tfh cells during the recall response. To address this hypothesis, LCMV GP-specific memory CD4+ T cells were generated in mice by acute LCMV infection that were treated with either DAC or PBS at 20 hours post infection. Thirty days later, mice were intranasally-infected with a recombinant strain of influenza PR8 that contains the GP^61-80^ epitope within the HA molecule (Sircy et al., 2024). We analyzed the recall response in the spleen, mLNs, and lung tissues at 7 days post-secondary infection (Fig. 2M). DAC treated recipients had more GP-specific secondary (2°) effector T cells in the mLNs but not in the spleen (Fig. 2N). Analysis of the Th1 cell response revealed that 2° Th1 cell cell numbers were unchanged in the lymphoid tissues of DAC treated mice (Fig. S1I-J); however, DAC treatment resulted in increased Tbet expression in 2° Th1 and Tfh cells (Fig. S1K). In the lung, we found a trending increase in GP66-77 tetramer+ CD4+ T cells (p=0.0658) (Fig. S1L) and a significant increase in IFNγ+TNFα+IL-2+ triple-producing cells in DAC treated recipients (Fig. S1M). This correlated with an increase in Granzyme B expression in DAC treated 2° Th1 cells (Fig. S1N). These data suggest that early DAC treatment during T cell priming enhanced the functionality of 2° Th1 cells.

Analysis of the Tfh response revealed an increase in DAC treated 2° Bcl6^HI^ GC Tfh cells within the lymphoid tissues (Fig. 2O-P) and increased Bcl6 expression on 2° Tfh cells (Fig. 2Q). In addition, we found that DAC treated recipients had significantly increased GC B cell numbers in the mLNs and spleen, and Hemagglutinin-specific GC B cells in the spleen (Fig. 2R). Collectively, these results demonstrate that early DAC treatment during T cell priming enhances germinal center Tfh cell differentiation and GC B cell support during viral rechallenge.

### Decitabine treatment during T cell priming impairs the lineage commitment of memory Th1 cells

Several pieces of data suggest that DAC treatment may be interfering with memory CD4+ T cell lineage commitment. First, we demonstrated that Th1 and Tfh cells acquire distinct *de novo* methylation programs which may be impacted by DAC treatment (Fig. 1). Second, early treatment with the methyltransferase inhibitor decitabine enhanced the GC Tfh cell response during primary and secondary viral challenge (Fig, 2). Phenotypically, DAC treated SMARTA cells expressed more Bcl6 at effector timepoints (Fig. 2F), and less Tbet at memory timepoints (Fig. 2L). Therefore, we hypothesized that early DAC treatment during priming impairs the lineage commitment of memory Th1 cells. To test our hypothesis, we sorted SMARTA memory cells from DAC- or PBS-treated recipients into CXCR5+ Ly6C^LO^ mTfh and CXCR5-Ly6C^HI^ mTh1 populations. These populations were then independently transferred into naïve B6 mice followed by LCMV infection of recipients (Fig. 3A-B). Seven days later, secondary SMARTA expansion was similar between DAC- and PBS-treated groups (Fig. 3C). In agreement with previous literature, we found that mTh1 cells that were generated in PBS-treated mice were lineage committed (Hale et al., 2013, Künzli et al., 2020). The majority of mTh1 progeny cells upregulated granzyme B and took on a Th1-like phenotype (Fig. 3D-E). Additionally, mTfh cells that were generated in PBS-treated mice produced more 2° GC Tfh cells relative to mTh1 cells (Fig. 3F-G). Thus, we analyzed the effects of early DAC treatment on the mTh1 recall response. DAC treated mTh1 cells produced significantly fewer granzyme B+ Th1 cells by percent (Fig. 3D-E), and significantly more 2° GC Tfh cells by both percent and number (Fig. 3F-G). Importantly, DAC treated mTh1 cells produced 2° Th1 cells that expressed less Tbet and aberrantly upregulated Tcf1 (Fig. 3H). Additionally, DAC treatment enhanced the ability of mTh1 cells to support B cell responses. Recipients of DAC-treated mTh1 cells had a significant increase in the numbers of plasmablasts and GC B cells (Fig. 3I-J). Overall, these data demonstrate that DAC treatment during T cell priming impairs the lineage commitment and function of mTh1 progeny cells during the memory recall response.

**Figure 3.**
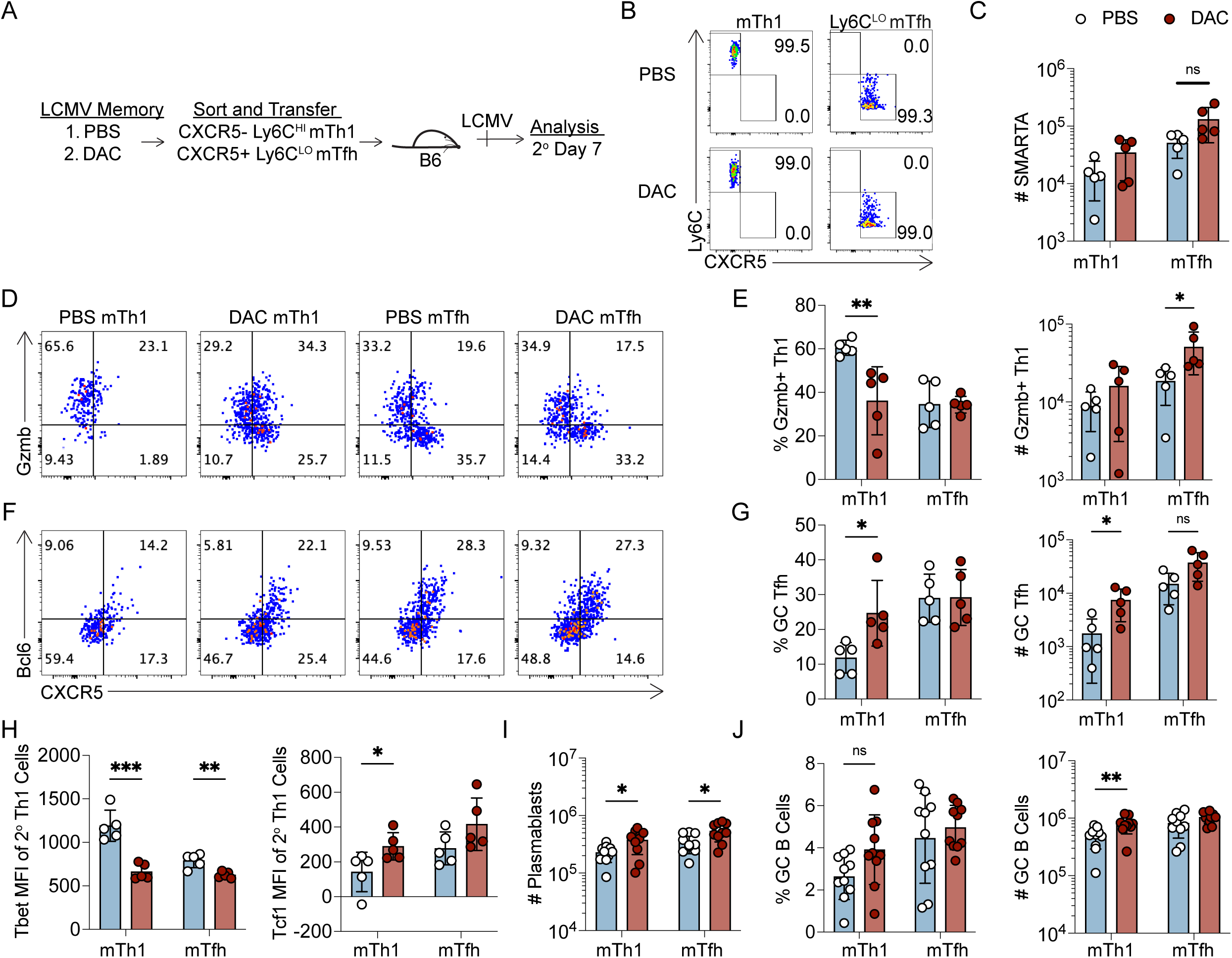
Decitabine treatment during T cell priming impairs the lineage commitment of memory Th1 cells. DAC or PBS treated SMARTA memory cells were sorted, and populations were independently transferred into naïve B6 mice which were then infected with LCMV. SMARTA T cell memory recall responses were analyzed 7 days post infection in the spleen. Data are representative of two independent experiments. B cell data are pooled from two experiments. (**A**) Experimental schematic. (**B**) FACS purity analysis of memory T cell populations post sort. (**C**) Number of SMARTA cells in the spleens. (**D**) Representative FACS plots of Granzyme B (Gzmb) and CXCR5 staining for Th1 identification, gated on SMARTA T cells. (**E**) Percent and Number of CXCR5-Gzmb+ Th1 cells. (**F**) FACS plots of Bcl6 and CXCR5 for GC Tfh identification, gated on SMARTA T cells. (**G**) Percent and number of CXCR5+ Bcl6^hi^ GC Tfh cells. (**H**) Tbet and Tcf1 MFIs of CXCR5-Th1 SMARTA cells. (**I**) Number of Plasmablasts. (**J**) Percent and number of GC B cells. Statistical significance was determined by an unpaired t test (* p<0.05, **p<0.01, ***p<0.001).

In comparison to the mTh1 cells, DAC treatment had a less remarkable effect on mTfh cell function and lineage commitment. DAC treated mTfh cells produced a similar frequency and number of GC Tfh cells (Fig. 3F-G) and GC B cell accumulation was unaffected by DAC treatment (Fig. 3K). In addition, while there was an increase in the number of 2° Th1 cells in recipients of DAC treated mTfh cells (Fig. 3D-E), these cells expressed less Tbet (Fig. 3H). Thus, DAC treatment early during priming had only a minor impact on the phenotype and function of mTfh progeny cells during the memory recall response.

### Dnmt3a restricts GC Tfh cell differentiation in a cell-intrinsic manner

Thus far, our data demonstrates loss of memory Th1 cell lineage commitment following early DAC treatment (Fig. 3), which may be due to inhibition of lineage-specific *de novo* methylation at *Tcf7* and other Tfh-associated genes (Fig. 1). Previous studies demonstrate that Dnmt3a is the predominantly upregulated *de novo* methyltransferase in activated CD4+ T cells (Gamper et al., 2009, Thomas et al., 2012). Thus, we hypothesized that DAC treatment may impair mTh1 lineage commitment by inhibiting Dnmt3a-mediated *de novo* methylation programing. To address this question, we first generated *Dnmt3a^flox/flox^* x *ERT2-cre* x SMARTA mice (CD45.1+). Donor *ERT2-cre* x SMARTA mice (either *Dnmt3a*^+/+^ or *^flox/flox^*) were treated with tamoxifen to induce Cre-mediated recombination. Naïve SMARTA cells from either WT or *Dnmt3a* conditional knockout (cKO) donor mice were adoptively transferred into B6 recipients that were then infected with LCMV. The CD4+ T cell response was analyzed 7 days later in the spleen. *Dnmt3a* deletion was validated by flow cytometry and qPCR (Fig. S2A-B). *Dnmt3a* deletion had little effect on SMARTA expansion in the spleen (Fig. 4A) and Tfh/Th1 differentiation was unaltered (Fig. 4B). There was also no difference in the number of IFNγ+ TNFα+ and IL-2+ triple producing SMARTA cells between the two groups (Fig. S2C). However, like in the DAC model, we found that *Dnmt3a* cKO SMARTA cells had a significant increase in the percent and number of Bcl6^HI^ GC Tfh cells (Fig. 4C-D) and Tfh cells expressed more Bcl6 (Fig. 4E). To test whether the increase in GC Tfh cells was due to a cell intrinsic effect, we co-transferred equal numbers of WT and *Dnmt3a* cKO SMARTA cells into B6 mice and infected with LCMV. While there was no difference in the percent of Tfh cells, *Dnmt3a* cKO SMARTA cells exhibited a significant increase in the percent of GC Tfh cells at 7 days post infection (Fig. 4F-G). These experiments demonstrate that Dnmt3a-dependent intrinsic programing restricts GC Tfh differentiation during viral infection.

**Figure 4.**
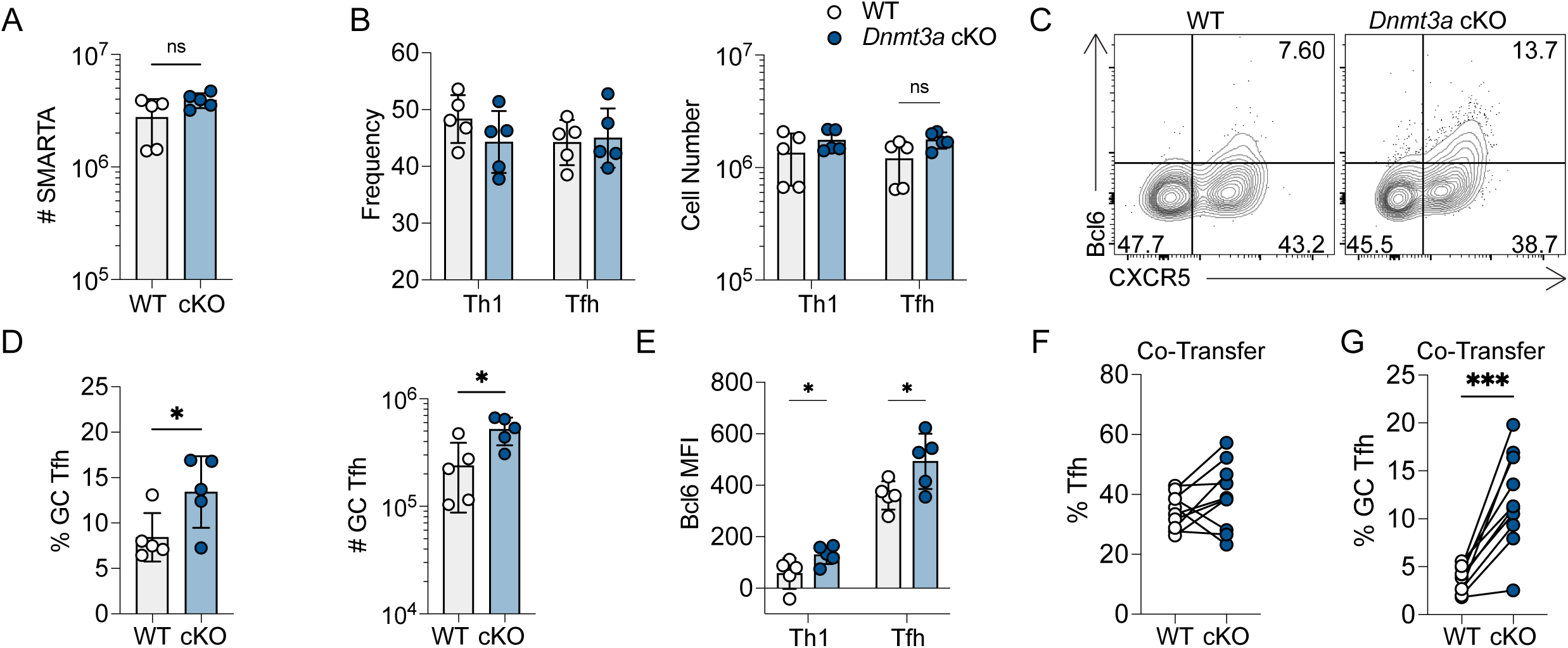
Dnmt3a restricts GC Tfh cell differentiation in a cell-intrinsic manner. (A-E) *ERT2-cre* x SMARTA (WT) or *Dnmt3a^flox/flox^* x *ERT2-cre* x SMARTA (cKO) mice were treated with tamoxifen to induce Dnmt3a deletion. SMARTA cells were transferred into B6 mice and were infected with LCMV. The CD4+ T cell responses were analyzed 7 days post infection within the spleen. Data are representative of three independent experiments. (**A**) Number of SMARTA T cells. (**B**) Percent and number of CXCR5+ Tfh or CXCR5-Th1 SMARTA cells. (**C**) Representative FACS plots of Bcl6 and CXCR5, gated on SMARTA T cells. (**D**) Percent and number of CXCR5+ Bcl6^HI^ GC Tfh SMARTA cells. (**E**) Bcl6 MFI of Tfh and Th1 SMARTA cells. Statistical significance was determined by an unpaired t test (* p<0.05, **p<0.01, ***p<0.001). (F-G) WT or KO SMARTA mice were treated with tamoxifen to induce *Dnmt3a* deletion. Mice were mixed at a 1 to 1 ratio and subsequently transferred into B6 mice. Mice were infected with LCMV, and CD4+ T cell responses were analyzed 7 days post infection. Data are representative of a single experiment. (**F**) percent CXCR5+ Tfh and (**G**) CXCR5+ Bcl6^HI^ GC Tfh SMARTA cells. Statistical significance was determined by a paired t test (* p<0.05, **p<0.01, ***p<0.001).

### Dnmt3a is dispensable for memory cell formation

We next analyzed the effect of *Dnmt3a* deficiency on memory formation. In initial adoptive transfer experiments, which were carried out using B6 mice as recipients, WT and *Dnmt3a* cKO SMARTA cells were detectable at effector timepoints but undetectable at memory timepoints (Fig. S2D-E). This issue was resolved by transferring WT and cKO SMARTA cells into Cre+ B6 recipients (Fig. S2D-E). Thus, we transferred either WT or cKO SMARTA cells into Cre+ B6 recipients, infected with LCMV, and analyzed the memory response in the spleen at 60+ days post infection. Loss of *Dnmt3a* had no effect on the number of SMARTA memory cells (Fig. S2F). Phenotypically, we observed a decrease in the frequency but not the number of CXCR5+ Ly6C^HI^ cells (Fig. S2G). In addition, cKO Ly6C^HI^ mTfh and Ly6C^LO^ mTfh cells expressed less Tcf1 and cKO mTh1 cells expressed less Tbet (Fig. S2H-I). Therefore, while *Dnmt3a* loss did not alter memory formation, it did reduce expression of Tfh and Th1 transcription factors at memory timepoints, suggesting impaired lineage commitment of *Dnmt3a* cKO mTfh and mTh1 cells.

### Dnmt3a limits plasticity and preserves functionality of memory Tfh and Th1 cells

To address the effects of *Dnmt3a* deletion on lineage commitment of memory T cells, WT or *Dnmt3a* cKO SMARTA memory cells were FACS sorted into mTh1, Ly6C^HI^ mTfh and Ly6C^LO^ mTfh subsets (Fig. S3A), and transferred into B6 recipients that were then infected with LCMV. The spleen and lung tissues were harvested at 7 days later (Fig. 5A). Across three repeat experiments, we observed an inconsistent reduction in the number of cKO SMARTA cells in the spleen (Fig. S3B). Next, we analyzed the effect of *Dnmt3a* deletion on the mTh1 recall response. *Dnmt3a* cKO mTh1 cells produced significantly fewer Tbet^HI^ 2° Th1 cells (Fig. 5B, E), and more Tcf1+ 2° Th1 cells (Fig. 5C, F). This correlated with reduced Tbet, IFNγ, and granzyme B expression in the spleen as well as reduced granzyme B expression in the lung (Fig. S3C-F). We next analyzed the plasticity of *Dnmt3a* cKO mTh1 cells and found a significant increase in CXCR5+ 2° Tfh cells (Fig. 5D, G). Despite this increase in 2° Tfh cells there was no increase in Bcl6^HI^ 2° GC Tfh cells generated from cKO mTh1 cells (Fig. 5D, H), and 2° Tfh cells from cKO mTh1 cells expressed less Bcl6 (Fig. 5I). These data demonstrate that Dnmt3a enhances the functionality of mTh1 progeny cells and suggests that Dnmt3a is required to silence the Tfh transcriptional program in mTh1 cells.

**Figure 5.**
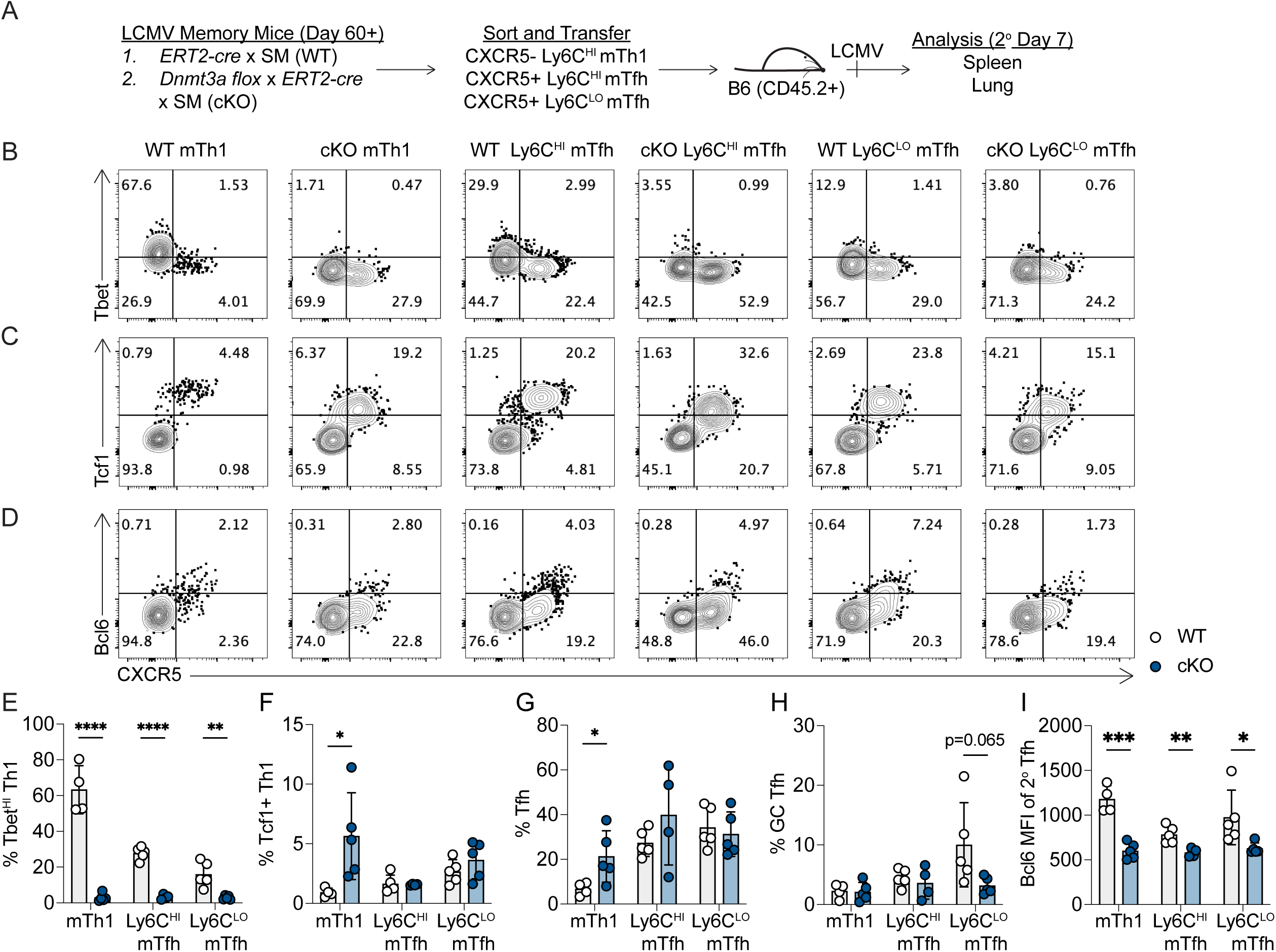
Dnmt3a limits plasticity and preserves functionality of memory Tfh and Th1 cells. (A-I) *ERT2-cre* x SMARTA (WT) or *Dnmt3a^flox/flox^* x *ERT2-cre* x SMARTA (cKO) mice were treated with tamoxifen to induce Dnmt3a deletion. SMARTA cells were transferred into B6 mice and were infected with LCMV. At 60+ days post infection, SMARTA cells were sorted from spleens into CXCR5-Ly6C^HI^ mTh1, CXCR5+ Ly6C^HI^ mTfh, and CXCR5+ Ly6C^LO^ mTfh, independently transferred into naïve B6 recipients, and mice were infected with LCMV. The recall analysis was analyzed in the spleen and lung at 7 days post-secondary infection. Data are representative of three independent experiments. (**A**) Experimental schematic. Representative FACS plots of (**B**) Tbet by CXCR5, (**C**) Tcf1 by CXCR5, and (**D**) Bcl6 by CXCR5. All FACS plots are gated on SMARTA cells. Quantification of the percent (**E**) CXCR5-Tbet^HI^ Th1 cells, (**F**) Tcf1+ Th1 cells, (**G**) CXCR5+ Tfh cells, and (**H**) CXCR5+ Bcl6^HI^ GC Tfh cells. (**I**) Quantification of Bcl6 MFI of CXCR5+ Tfh cells. Statistical significance was determined by an unpaired t test (* p<0.05, **p<0.01, ***p<0.001).

We then analyzed the effect of *Dnmt3a* deletion on the mTfh recall response. While *Dnmt3a* cKO Ly6C^LO^ mTfh cells produced similar frequencies of CXCR5+ 2° Tfh cells (Fig. 5G), there was a trending reduction in Bcl6^HI^ 2° GC Tfh cells (p=0.065) (Fig. 5D, H) and 2° Tfh cells expressed significantly less Bcl6 and Tcf1 (Fig. 5I, S3G). This impairment in the functional Tfh response from the *Dnmt3a* cKO mTfh subsets was not due to enhanced plasticity. To the contrary, *Dnmt3a* cKO Ly6C^LO^ mTfh and Ly6C^HI^ mTfh cells also produced significantly fewer Tbet^HI^ 2° Th1 cells (Fig. 5B, E). Together, these findings indicate that Dnmt3a is essential for mTfh functionality while having little effect on mTfh plasticity, suggesting that Dnmt3a is required to silence genes that repress *Bcl6* to enable GC Tfh differentiation during the memory recall response.

We used an additional Cre model system (*Dnmt3a^flox/flox^* x *CD4-Cre* x SMARTA) to validate our results. We FACS sorted *CD4-Cre* x SMARTA memory cells that were either *Dnmt3a^+/+^* (WT) or *Dnmt3a ^flox/flox^* (KO), independently transferred them into B6 recipients, and infected with LCMV (Fig. S3H). In agreement with the ERT2-Cre model, recipients of KO memory SMARTA cells expanded normally, had a trending reduction in 2° GC Tfh cells, and KO 2° Tfh cells expressed significantly less Bcl6 (Fig. S3I-K). Importantly, we found a significant reduction in GC B cells by percent and a trending reduction by number (p=0.075) (Fig. S3L). We also found evidence of impaired Th1 cell responses and KO 2° Th1 cells aberrantly upregulated Tcf1 (Fig. SM-O). Therefore, we concluded that *Dnmt3a* deficient memory T cells (*ERT2-Cre* and *CD4-Cre*) had substantial impairments in function during the memory recall response to viral infection, suggesting that Dnmt3a is required to silence alternative T helper lineages in order to acquire specialized effector functions.

### Dnmt3a silences genes associated with alternative T helper lineages in Tfh and Th1 cells

*Dnmt3a* deficiency impaired the lineage commitment and functionality of mTh1 cells, which we hypothesized was due to loss of *de novo* methylation programing at Tfh-associated genes. In addition, *Dnmt3a* deficiency impaired the functionality of mTfh cells, and we hypothesized that this was due to loss of *de novo* methylation programing at genes encoding repressors of *Bcl6*. To determine whether Dnmt3a was necessary for *de novo* methylation at genes associated with alternative T helper lineages, we performed whole genome enzymatic methylation sequencing (WGEM-seq) on WT and *Dnmt3a* KO (*CD4-Cre*) Tfh and Th1 cells sorted from LCMV infected mice at 7 days post infection. DMRs were identified between WT and KO cells of the same lineage. This analysis revealed 26,513 DMRs in Tfh cells and 27,937 DMRs in Th1 cells (Fig. 6A). The majority of DMRs were hypomethylated in KO compared to WT and occurred within intragenic regions (Fig. 6A, S4A). Dnmt3a-dependent DMRs were associated with enrichment of H3K4me1 peaks (in Naïve CD4+ T cells) as well as ETS and RUNX motifs, which is consistent with T cell regulatory regions (Fig. S4B-C) (Placek et al., 2017, Zhong et al., 2022). Importantly, most *de novo* DMRs were lost in KO Tfh and Th1 cells (Fig. S4D). These data indicate that Dnmt3a is the predominant *de novo* methyltransferase in Tfh and Th1 cells and that it is involved in silencing T cell specific regulatory regions.

**Figure 6.**
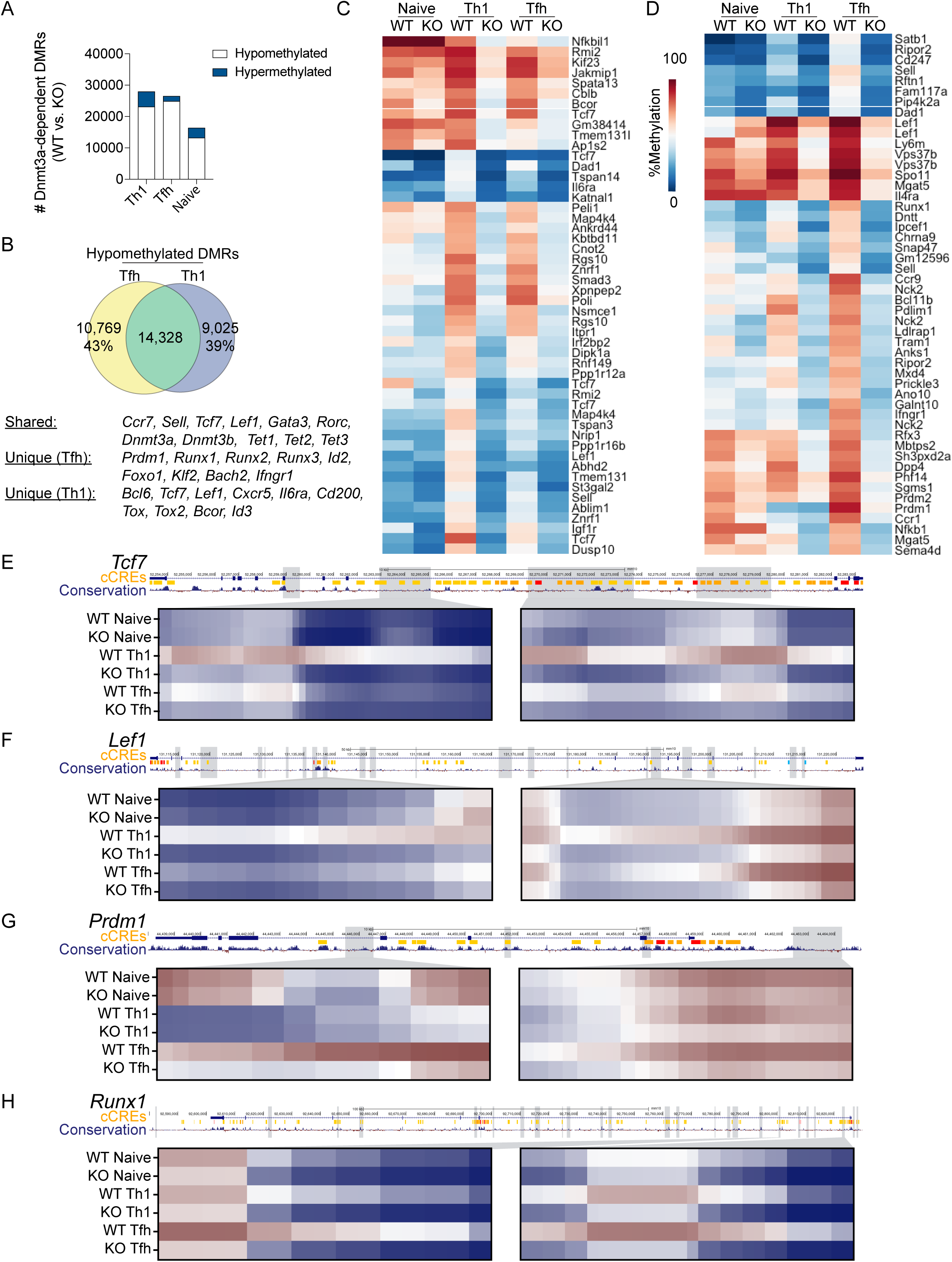
Dnmt3a silences genes associated with alternative T helper lineages in Tfh and Th1 cells. Whole genome enzymatic methylation sequencing was performed on *CD4-cre* x SMARTA (WT) and *Dnmt3a^flox/flox^* x *CD4-cre* x SMARTA (KO) Tfh and Th1 cells sorted at 7 days post LCMV infection. WT and KO naïve CD4+ T cells were also analyzed as controls. Differentially methylated regions (DMRs) were called between WT and KO cells of the same lineage. Data are representative of three biological replicates. (**A**) Quantification of the total number of DMRs per cell type (WT vs KO). Hypermethylated regions are blue and hypomethylated regions are white. (**B**) Venn diagram of overlapping hypomethylated DMRs in Tfh and Th1 cells. Heatmaps of (**C**) the top 50 Th1-specific DMRs and (**D**) the top 50 Tfh-specific DMRs. DMRs were ranked by mean difference between WT Tfh and WT Th1 cells. UCSC genome browser snapshots of (**E**) *Tcf7*, (**F**) *Lef1*, (**G**) *Prdm1*, and (**H**) *Runx1*. We also show tracks for ENCODE identified cis-candidate regulatory elements (cCREs) and mammalian conservation. Grey boxed identify hypomethylated DMRs. In the heatmaps below, select DMRs are shown. Each column represents a CpG site across the DMR. Significant DMRs were identified using BSmooth DMR finder. DMRs have at least 10 CpGs and an absolute mean difference of >0.1.

Our previous analysis of *de novo* methylated regions demonstrated extensive programing that was shared between Tfh and Th1 cells (Fig. 1B-D). In agreement with this, we found that the majority of Th1 and Tfh DMRs were shared (Fig. 6B). This included DMRs at genes associated with memory cells (*Ccr7, Sell, Tcf7, Lef1, Id3*), Th2/Th17 differentiation (*Gata3, Rorc*), and DNA methylation programming (*Dnmt3a, Dnmt3b, Tet2*). We also found extensive lineage-specific methylation (Fig. 6B-D). Heatmap analysis of the top 50 Th1-specific DMRs included several genes which are involved in Tfh differentiation, including

*Tcf*7, *Lef1, Il6ra* and *Bcor* (Fig. 6C). In Tfh cells, the top 50 DMRs included several genes which are involved in Th1 differentiation, including *Prdm1, *Runx1*,* and *Ifngr1* (Fig. 6D). We further analyzed select Th1 DMRs at the *Tcf7, Lef1* and *Bcl6* loci, revealing extensive *de novo* methylation at intronic regions with high-degrees of mammalian conservation (Fig. 6E-F, S4E). To our surprise, the first intron of *Bcl6* was more methylated in Tfh cells (Fig. S4E). Additionally, motif analysis found specific enrichment of Tcf1 motifs within 24% of Dnmt3a DMRs in Th1 cells (Fig. S4C). For Tfh cells, we chose to further analyze the genes *Prdm1, *Runx1*,* and *Runx2,* revealing extensive *de novo* methylation that occurred at intronic regions with a high degree of mammalian conservation (Fig. 6G-H, S4F). Motif analysis found specific enrichment of *Runx1* motifs within 23% of Dnmt3a DMRs in Tfh cells (Fig. S4C). Together, these data demonstrate that Dnmt3a is important for *de novo* methylating genes associated with alternative lineages.

It was unexpected that *de novo* methylation in the first intron of *Bcl6* was stronger in Tfh cells compared to Th1 cells (Fig. S4E), suggesting that DNA methylation in this region may promote *Bcl6* expression. In agreement with this idea, one publication analyzed DNA methylation in human GC B cells and plasma cells and found that GC B cells more strongly methylated the first intron of *Bcl6* (Lai et al., 2010)*. Lai et al.* argue that DNA methylation within this region inhibits binding and repression of *Bcl6* by CTCF, thereby promoting *Bcl6* expression in GC B cells (Lai et al., 2010). Thus, we first asked whether our intronic DMRs overlapped with the human CpG islands interrogated in their study using LiftOver, and we found that three of the human CpG islands overlapped with two Dnmt3a-dependent DMRs identified in our study (Fig. S4E). Next, using publicly available CTCF ChIP-seq data (SRX5086069) from murine naïve CD4+ T cells (Pham et al., 2019), we found evidence of two CTCF peaks overlapping with two Dnmt3a-dependent DMRs (Fig. S4E). Therefore, the impairment in Bcl6 expression in *Dnmt3a* deficient 2° Tfh cells may not only be due to failure to silence repressors of *Bcl6,* but also due to hypomethylation of a *Bcl6* silencer region.

To determine genes where Dnmt3a may silence expression via *de novo* methylation in primary effector cells, we performed RNA sequencing on Tfh and Th1 cells sorted from LCMV infected mice at 7 days post infection. Differentially expressed genes (DEGs) were identified between WT and *Dnmt3a* KO cells of the same cell type. We found 1126 Tfh (WT vs KO) and 58 Th1 DEGs (WT vs KO) (Fig. S4G). We identified genes that were upregulated and hypomethylated in cKO Th1 cells. This included the Tfh-associated gene *Lef1*, which was more expressed by transcript and protein levels (Fig. S4H-I). *Dnmt3a* KO Tfh cells upregulated and hypomethylated several Th1-associated genes, including *Runx1* which was more highly expressed by transcript and protein levels compared to WT cells (Fig. S4J-K). Thus, Dnmt3a-dependent methylation silences genes associated with alternative T helper lineages including *Lef1* in Th1 cells and *Runx1* in Tfh cells during the primary response.

### Early decitabine treatment partially inhibits Dnmt3a-mediated methylation programming at Tfh-associated genes

Our analysis of the methylomes of knockout cells demonstrates that Dnmt3a is required for silencing genes associated with alternative T helper lineages. DAC treatment and *Dnmt3a* deletion resulted in memory Th1 cells that exhibited impaired lineage commitment during the recall response, suggesting that DAC inhibited Dnmt3a-mediated silencing of the Tfh program in Th1 cells. However, DAC treatment and *Dnmt3a* deletion had divergent effects on the functionality of memory T cells, especially regarding 2° GC Tfh cell formation, suggesting that the mechanism by which DAC treatment enhances memory T cell plasticity and functionality may be more complex. Mechanistically, decitabine treatment leads to Dnmt3a degradation (Christman, 2002, Derissen et al., 2013, Seelan et al., 2018). To address whether Dnmt3a was inhibited in DAC treated SMARTA cells, we measured Dnmt3a levels in early activated SMARTA cells. Naïve SMARTA cells were transferred into B6 mice, infected with LCMV, and treated with DAC 20 hours later. At 3 days post infection, we found a small but significant decrease in the MFI of Dnmt3a within DAC treated SMARTA cells (Fig. 7A). These data demonstrate that DAC treatment led to Dnmt3a degradation within SMARTA cells, although they may suggest that DAC mediated inhibition of Dnmt3a is temporary and incomplete.

**Figure 7.**
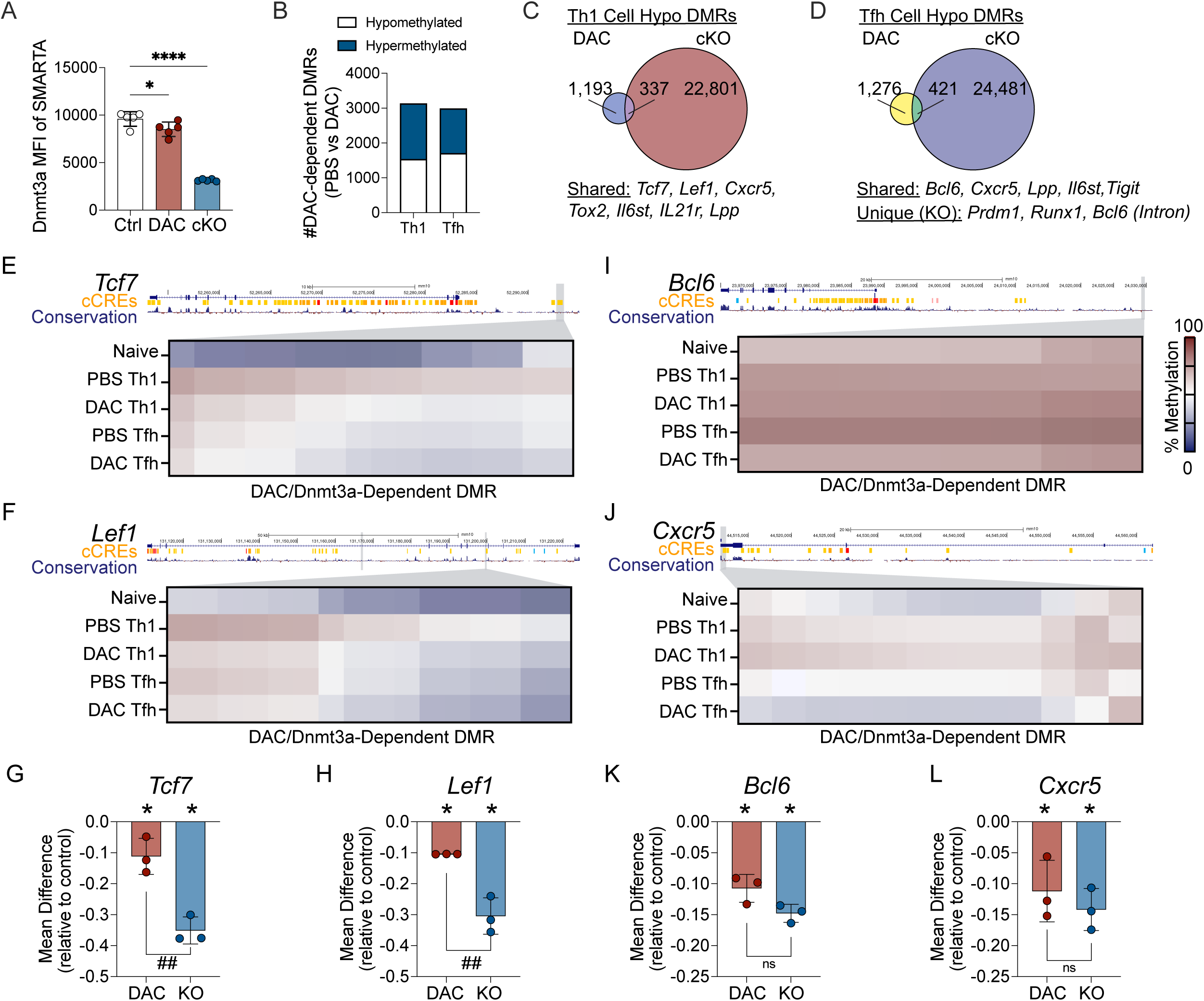
Early decitabine treatment partially inhibits Dnmt3a-mediated methylation programing at Tfh-associated genes. (A) *ERT2-cre* x SMARTA (WT) or *Dnmt3a^flox/flox^* x *ERT2-cre* x SMARTA (cKO) mice were treated with tamoxifen to induce Dnmt3a deletion. Cells were adoptively transferred into B6 recipients and intravenously infected with LCMV. WT cells received either PBS (WT-PBS) or DAC (WT-DAC) while all cKO cells received PBS (cKO-PBS). At 3 days post infection, the T cell response was analyzed in the spleen. Data are representative of a single experiment. (**A**) The Dnmt3a MFI of WT, cKO or DAC treated SMARTA T cells is shown in the figure. Statistical significance was determined by an unpaired t test (* p<0.05, **p<0.01, ***p<0.001). (B-L) Whole genome enzymatic methylation sequencing was performed on PBS or DAC treated Tfh and Th1 cells sorted at 7 days post LCMV infection. Differentially methylated regions (DMRs) were called between PBS and DAC treated cells of the same lineage. Data are representative of three biological replicates. (**B**) Quantification of the number and directionality of DMRs. For C and D, overlap analysis was performed on DAC-dependent hypomethylated DMRs and Dnmt3a-dependent hypomethylated DMRs identified from (**C**) Th1 or (**D**) Tfh cells. (E-F) UCSC genome browser snapshots of Th1 cell DMRs at (**E**) *Tcf7*, and (**F**) *Lef1*. All shown DMRs are DAC-dependent DMRs (PBS Th1 vs DAC Th1) that overlap with Dnmt3a-dependent DMRs (WT Th1 vs KO Th1). (G-H) Mean difference in methylation is shown across the DAC-dependent Th1 DMR at (**G**) *Tcf7* and (**H**) *Lef1*. For I and J, UCSC genome browser snapshots of Tfh cell DMRs at (**I**) *Bcl6*, and (**J**) *Cxcr5*. All shown DMRs are DAC-dependent DMRs (PBS Tfh vs DAC Tfh) that overlap with Dnmt3a-dependent DMRs (WT Tfh vs KO Tfh). For K and L, the mean difference in methylation is shown across the DAC-dependent Tfh DMR at (**K**) *Bcl6* and (**L**) *Cxcr5*. For plots comparing KO and DAC, * symbol represents significant DMR. # symbol represents significant difference between groups as determined by a t test. For all UCSC genome browser snapshots, we also show tracks for ENCODE identified cis-candidate regulatory elements (cCREs) and mammalian conservation. Grey boxed show DMRs and select DMRs are highlighted below. Each column represents a CpG site across the region. Significant DMRs were identified using BSmooth DMR finder. DMRs have at least 10 CpGs and an absolute mean difference of >0.1.

To determine if DAC treatment impaired Dnmt3a-mediated methylation programing, we performed WGEM-seq on Tfh and Th1 cells from PBS or DAC treated mice at 7 days post infection. DMRs were identified between PBS and DAC treated cells of the same lineage. This analysis revealed 2984 Tfh DMRs and 3127 Th1 DMRs that were DAC-dependent (Fig. 7B). Approximately half of the DMRs were hypermethylated (Fig. 7B), suggesting that DAC treatment may have indirect effects on the acquisition of DNA methylation programing. To determine if DAC treatment resulted in hypomethylation of Dnmt3a-dependent DMRs, we identified overlapping (shared) DMRs. This analysis identified 337 Th1 DMRs and 421 Tfh DMRs that were hypomethylated in both the DAC treatment and *Dnmt3a* KO models, suggesting that DAC inhibited Dnmt3a-mediated methylation programing at several hundred regions (Fig. 7C-D). Importantly, DAC treatment failed to inhibit Dnmt3a-mediated methylation at 22,901 regions in Th1 cells and 24,481 regions in Tfh cells (Fig. 7C-D), supporting the conclusion that early DAC treatment temporarily inhibited Dnmt3a-mediated programing in Tfh and Th1 cells.

Analysis of shared DMRs between *Dnmt3a* cKO and DAC treated Th1 cells revealed hypomethylation of several Tfh genes, including *Tcf7, Lef1, Cxcr5, Tox2,* and *Il6st* (Fig. 7C). At *Tcf7,* DAC inhibited Dnmt3a-dependent methylation at an upstream region that overlapped with a candidate cis regulatory element (cCRE) (Fig. 7E). At *Lef1,* DAC treatment also inhibited Dnmt3a-dependent *de novo* methylation at two intronic regions, including one that overlapped with a cCRE (Fig. 7F). Next, we directly compared the effect of *Dnmt3a* deletion and DAC treatment on the acquisition of *de novo* methylation at *Tcf7* and *Lef1*. This analysis demonstrated that DAC treatment only partially inhibited the acquisition of Dnmt3a-dependent *de novo* methylation (Fig. 7G-H). Thus, decitabine treatment during T cell priming partially and temporarily inhibited Dnmt3a-dependent *de novo* methylation at Tfh-associated genes in Th1 cells, impairing memory Th1 cell lineage commitment.

Analysis of shared DMRs between *Dnmt3a* cKO and DAC treated Tfh cells revealed hypomethylation of several Tfh genes within Tfh cells, including *Bcl6, Cxcr5, Lpp* and *Il6st* (Fig. 7D). DAC treatment inhibited Dnmt3a-mediated *de novo* methylation at a region upstream of *Bcl6* as well as within the gene *Cxcr5* (Fig. 7I-J). Inhibition of Dnmt3a-mediated *de novo* methylation at these two loci were nearly complete (Fig. 7K-L). These data suggest that early Dnmt3a-mediated *de novo* methylation directly represses Bcl6 expression and GC Tfh cell differentiation in CD4+ T cells during primary viral infection.

### Dnmt3a deficiency, but not early decitabine treatment, impairs silencing of the loci encoding Blimp1 and *Runx1*

Our methylation analyses demonstrate that early decitabine treatment inhibits Dnmt3a-mediated methylation programing at several Tfh-associated genes within Tfh and Th1 cells. However, these data fail to explain the differences we observed in 2° GC Tfh cell formation between the DAC and Dnmt3a KO models. Specifically, *Dnmt3a* KO (from CD4-Cre and ERT2-Cre models) memory Tfh cells expressed less Bcl6 (Figs. 5I, S3K), failed to differentiate into 2° GC Tfh cells (Figs. 5H), and were worse at supporting GC B cells during the recall response (Fig. S3L). On the other hand, decitabine treated memory T cells expressed more Bcl6 (Fig. 2Q), produced more 2° GC Tfh (Figs. 2P, 3G), and were better at supporting GC B cell formation during the recall response (Figs. 2R). Our DNA methylation analysis underscored that decitabine mediated inhibition of Dnmt3a was temporary and incomplete (Fig. 7C-D). Thus, we hypothesized that the differences in Bcl6 re-expression may be due to *Dnmt3a* KO Tfh cells failing to *de novo* methylate loci encoding repressors of Bcl6, which may properly acquire *de novo* methylated following DAC treatment. To address this question, we first analyzed the effects of decitabine treatment on Dnmt3a-mediated *de novo* methylation at several repressors of Tfh differentiation and function. At *Prdm1* and *Runx1*, we found that DAC treatment had no effect on the acquisition of Dnmt3a-mediated methylation at select DMRs (Fig. 8A-C). In addition, DAC treatment had little effect on the acquisition of Dnmt3a-mediated methylation at the silencer region within the first intron of *Bcl6* (Fig. S5A-B). Broadly, DAC treatment failed to inhibit Dnmt3a-mediated methylation at any DMRs associated with *Prdm1* (11 DMRs), *Runx1* (43 DMRs)*, Foxo1* (26 DMRs), *Runx2* (29 DMRs), *Runx3* (20 DMRs), and *Id2* (3 DMRs) (Figs. 10D, S5C). Overall, these data demonstrate that DAC treatment did not inhibit Dnmt3a-mediated methylation at several known repressors of Tfh differentiation and function.

**Figure 8.**
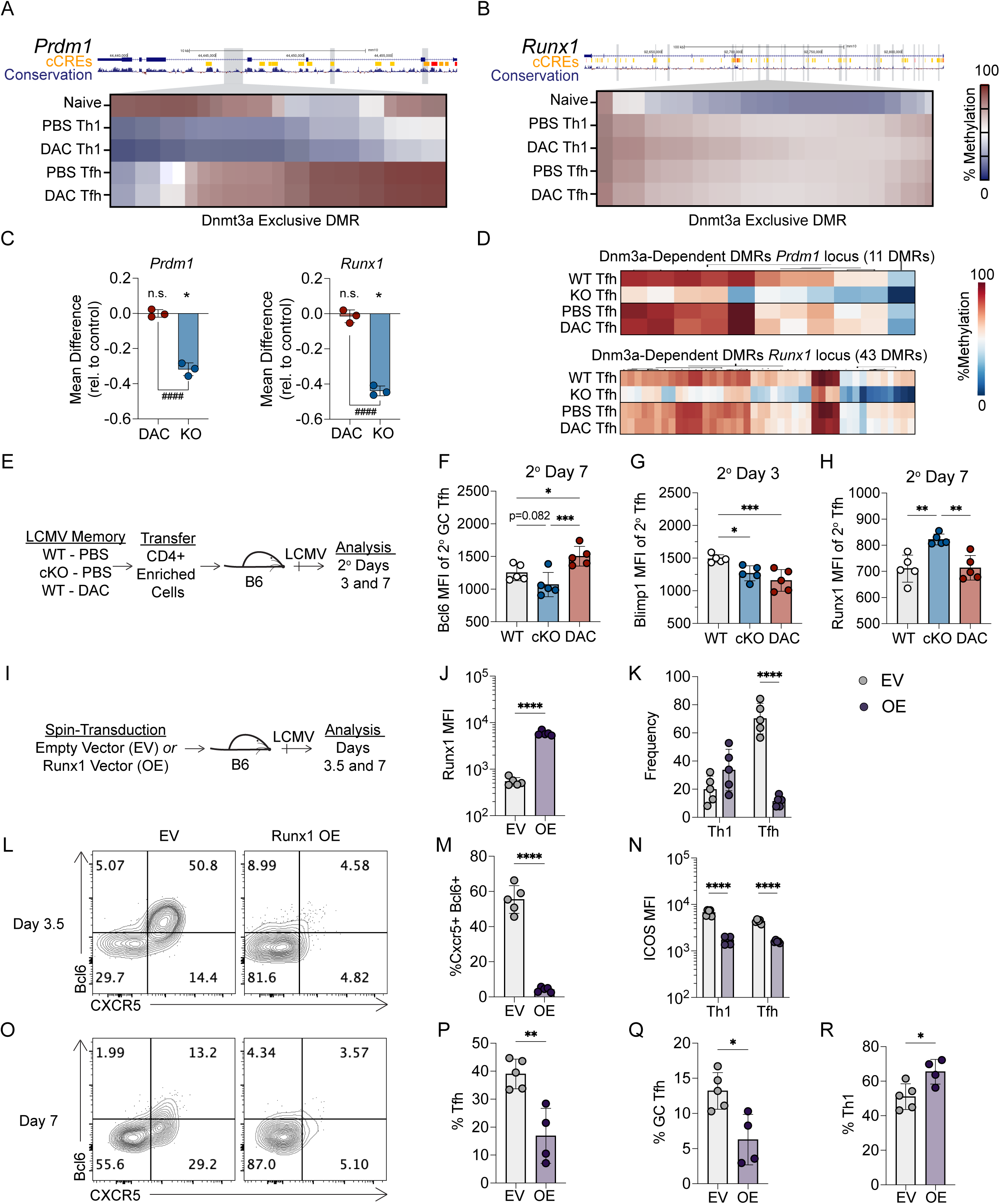
Dnmt3a deficiency, but not early decitabine treatment, impairs silencing of the loci encoding Blimp1 and *Runx1*. (A-D) Additional analysis of whole genome methylation sequencing data from figure 7. UCSC genome browser snapshots of Tfh cell Dnmt3a-dependent DMRs (WT Tfh vs KO Tfh) at (**A**) *Prdm1* and (**B**) *Runx1*. The shown DMRs are not DAC-dependent DMRs (PBS Tfh vs DAC Tfh). (**C**) The mean difference in methylation is shown across the Dnmt3a-dependent Tfh DMR at *Prdm1* and *Runx1.* (**D**) Mean methylation is shown for all Dnmt3a-dependent Tfh DMRs at the genes *Prdm1* and *Runx1*. For plots comparing KO and DAC, * symbol represents significant DMR. # symbol represents significant difference between groups as determined by a t test. (E-H) *ERT2-cre* x SMARTA (WT) or *Dnmt3a^flox/flox^* x *ERT2-cre* x SMARTA (cKO) mice were treated with tamoxifen to induce Cre-mediated deletion. Naïve CD4+ T cells were transferred into mice which were subsequently infected with LCMV. Mice were treated with either PBS or DAC at 20 hours post infection. At memory timepoint, CD4+ T cells were enriched from the spleen, and transferred into new naïve B6 mice. Mice were infected with LCMV and SMARTA cells were analyzed at 3 or 7 days post infection. Data are representative of two independent experiments. (**E**) Experimental schematic. (**F**) Bcl6 MFI of secondary GC Tfh at 7 days post infection, (**G**) Blimp1 MFI of secondary Tfh at 3 days post infection, or (**H**) *Runx1* MFI of secondary Tfh at 7 days post infection. Statistical significance was determined by an unpaired t test (* p<0.05, **p<0.01, ***p<0.001). (I-R) *Runx1* overexpression analysis of SMARTA T cells at 3.5 or 7 days post infection. Briefly, SMARTA cells were preactivated for 24 hours, then spin transduced with retroviral supernatant for 2 hours at 37 degrees. Cells were transferred into mice and infected with LCMV. Spleens were harvested for analysis. For intracellular staining, mCherry+ cells were sorted before intracellular staining. Data are representative of two independent experiments. (**I**) Experimental schematic. (**J**) *Runx1* MFI of mCherry+ SMARTA cells. (**K**) Quantification of Tcf1+ CXCR5+ Tfh or Tcf1-CXCR5-Th1 cells at 3.5 days post infection. (**L**) Representative FACS plots of CXCR5 and Bcl6 gated on mCherry+ SMARTA cells. (**M**) Quantification of CXCR5+ Bcl6+ Tfh cells at 3.5 days post infection. (**N**) Icos MFI of CXCR5+ CD25-Tfh and CXCR5-CD25+ Th1 cells at 3.5 days post infection. (**O**) Representative FACS plots of CXCR5 and Bcl6 gated on mCherry+ SMARTA cells. (**P**) Quantification of CXCR5+ Tfh, (**Q**) CXCR5+ Bcl6^HI^ GC Tfh, or (**R**) Tbet^HI^ Th1 cells at 7 days post infection. Statistical significance was determined by an unpaired t test (* p<0.05, **p<0.01, ***p<0.001).

To determine whether repressors of Tfh differentiation were aberrantly upregulated in *Dnmt3a* cKO 2° Tfh cells, LCMV derived SMARTA memory cells (Day 60) were generated that were either WT (PBS-treated), *Dnmt3a* cKO (PBS-treated), or WT (DAC-treated).

Memory CD4+ T cells from each group were enriched and transferred into B6 recipients that were subsequently infected with LCMV. We then analyzed the expression of transcription factors that repress Tfh differentiation at days 3 and 7 during the memory recall response (Fig. 8E). As anticipated, DAC treated 2° GC Tfh cells expressed more Bcl6 relative to WT (PBS treated) controls, while *Dnmt3a* cKO 2° GC Tfh cells had a trending reduction in Bcl6 relative to WT 2° GC Tfh cells (p=0.0852) and a significant reduction in Bcl6 relative to DAC treated 2° GC Tfh cells (Fig. 8F). To our surprise, both *Dnmt3a* cKO and DAC treated 2° Tfh cells downregulated Blimp1 at 3 days post infection (Fig. 8G), as well as Foxo1 and Runx3 (Fig. S5D). Id2 expression was trending lower in *Dnmt3a* cKO 2° Tfh, while Runx2 expression was unaltered (Fig. S5D). *Dnmt3a* cKO 2° Tfh cells did aberrantly upregulate *Runx1* while DAC treated 2° Tfh cells expressed normal levels of *Runx1* relative to the control (Fig. 8H). This correlated with reduced Icos expression in 2° GC Tfh cells (Fig. S5E), which is inhibited by Runx factors (Baessler et al., 2022, Choi et al., 2020). These data demonstrate that Dnmt3a is required to silence *Runx1* in 2° Tfh cells, and that early DAC treatment did not inhibit Dnmt3a-mediated silencing of *Runx1*.

Several studies suggest that *Runx1* may repress aspects of the Tfh transcriptional program (Wong et al., 2012, Hatzi et al., 2015, Baessler et al., 2022). Therefore, we next determined the effect of *Runx1* overexpression on GC Tfh differentiation and Bcl6 expression (Fig. 8I). *Runx1* overexpression was validated by flow cytometry (Fig. 8J).

Runx1 overexpressing SMARTA cells (OE) produced fewer Tcf1+ CXCR5+ and Bcl6+ Tfh cells by percent at 3.5 days post infection (Fig. 8K-M). This correlated with a strong reduction in Icos in both early Tfh and Th1 cells (Fig. 8N). At 7 days post infection, there were significantly fewer Tfh and Bcl6^HI^ GC Tfh cells (Fig. 8O-Q). In contrast, there were more Tbet^HI^ Th1 cells (Fig. 8R), demonstrating that *Runx1* represses GC Tfh cell formation and promotes Th1 cell formation. Overall, our data demonstrates that Dnmt3a is required to silence *Runx1* in Tfh cells to promote GC Tfh differentiation during the memory recall response. Furthermore, these findings indicate that DAC treatment inhibited Dnmt3a-mediated methylation at several Tfh genes (*Bcl6* and *Cxcr5*) but did not inhibit Dnmt3a-mediated methylation at genes encoding repressors of Tfh differentiation (*Prdm1* and *Runx1*), enhancing GC Tfh generation during the memory recall response.

## DISCUSSION

Acute viral infection generates Th1 and Tfh cells that give rise to distinct memory Th1 and Tfh subsets that recall their specialized effector functions upon rechallenge (Marshall et al., 2011, MacLeod et al., 2011, Morita et al., 2011, Pepper et al., 2011, Lüthje et al., 2012, Weber et al., 2012, Hale et al., 2013, Choi et al., 2013, Künzli et al., 2020, Baessler et al., 2023, Feng et al., 2024). Dnmt3a has been shown to play a role in effector T helper differentiation and commitment (Thomas et al., 2012, Gamper et al., 2009, Yu et al., 2012); however, whether Dnmt3a-dependent *de novo* methylation regulates the long-term lineage commitment and functionality of memory T cells was unclear. Recent work demonstrated the importance of DNA methylation programing, as Tet2-dependent DNA demethylation enforces the lineage commitment of memory Tfh and Th1 cells (Baessler et al., 2023). In this study, we demonstrate that the acquisition of *de novo* methylation programing during primary effector Tfh and Th1 differentiation is maintained in respective memory Tfh and Th1 cells, indicating that lineage-specific *de novo* methylation programing is foundational to the identity of memory Tfh and Th1 cells. In agreement with this idea, loss of *Dnmt3a* impaired the lineage commitment of memory Th1 cells. *Dnmt3a* deficiency also diminished the functionality of secondary Tfh and Th1 cells, and de-repressed several genes associated with the alternative T helper lineage. This included aberrant hypomethylation and expression of genes including *Tcf7* in secondary Th1 cells and *Runx1* in secondary Tfh cells.

Importantly, we demonstrate that *Runx1* overexpression suppressed GC Tfh cell differentiation, indicating that Dnmt3a is required to silence *Runx1* to enable secondary GC Tfh cell formation. Therefore, we conclude that Dnmt3a-dependent *de novo* methylation programing is required to enforce the lineage commitment of memory Th1 and Tfh cells by silencing key loci encoding transcription factors in order to preserve lineage-specific effector functions. Interestingly, treatment with the DNA methyltransferase inhibitor decitabine early during T cell priming increased memory Th1 cell plasticity, and in contrast to *Dnmt3a* deficient memory cells, resulted in enhanced generation of secondary GC Tfh cells. Early decitabine treatment partially inhibited the acquisition of Dnmt3a-medaited *de novo* methylation at several Tfh-associated genes in Tfh and Th1 cells, but importantly did not inhibit *de novo* methylation at *Runx1* and *Prdm1* in Tfh cells. Thus, our study demonstrates that partial inhibition of Dnmt3a-mediated *de novo* methylation using decitabine enhances memory Th1 cell plasticity and Tfh cell functionality.

We observed that *Dnmt3a* deletion enhanced memory Th1 cell plasticity, enabling some degree of secondary Tfh differentiation, albeit with impaired Bcl6 expression. Importantly, the *Bcl6* locus was more strongly methylated in Tfh cells at several intronic loci, suggesting a more complex relationship between Dnmt3a activity and regulation of Bcl6 expression. Th1 cells acquired lineage-specific *de novo* methylation at several intronic regions within the *Tcf7* and *Lef1* genes, and in line with this, Lef1 is upregulated in primary *Dnmt3a* deficient Th1 cells and Tcf1 is upregulated in *Dnmt3a* cKO secondary Th1 cells.

Previous research demonstrated that Tcf1 plays an early role in Tfh differentiation during viral infection by directly promoting *Bcl6* expression (Choi et al., 2015, Xu et al., 2015, Wu et al., 2015, Shao et al., 2019), suggesting that Dnmt3a indirectly silences *Bcl6* expression in Th1 cells. In addition, TCF/LEF motifs were enriched within approximately a quarter of Dnmt3a-dependent DMRs, suggesting that Dnmt3a may restrain accessibility to Tcf1 and Lef1 binding sites to further enforce Th1 lineage commitment. Secondary *Dnmt3a* cKO Th1 cells also have reduced expression of Tbet, IFNγ, and granzyme B relative to WT, and this may be attributed to Tcf1 repressing the Th1 effector program (Choi et al., 2015, Xu et al., 2015). Together, these findings support a model whereby Th1 cells require Dnmt3a-dependent *de novo* methylation to silence Tcf1- and Lef1-mediated transcriptional programing to enforce Th1 cell lineage commitment and enable Th1 effector functions (Fig. S5F).

Multiple groups have reported that memory Tfh cells induced by viral infection have higher plasticity relative to memory Th1 cells (Hale et al., 2013, Künzli et al., 2020, Baessler et al., 2023). Manipulation of Dnmt3a-dependent methylation programing was not without consequence however, and our data strongly implicates Dnmt3a-mediated programing during priming for preserving Tfh functionality during the memory recall response. We propose a model whereby Dnmt3a safeguards Tfh functionality by silencing several repressors of Tfh differentiation, including *Prdm1* and *Runx1* (Fig. S5F). Our data clearly show that Dnmt3a was required for *de novo* methylation at *Prdm1*, although we did not observe an increase in Blimp1 expression in *Dnmt3a* cKO Tfh cells at the timepoints analyzed. Importantly, Dnmt3a was required to methylate and repress *Runx1* expression in primary and secondary Tfh cells. Prior to our study, the evidence that *Runx1* repressed Tfh differentiation and function was correlative (Wong et al., 2012, Baessler et al., 2022). Here, we found that *Runx1* overexpression represses GC Tfh differentiation during viral infection, which is likely via repression of ICOS expression in early Tfh and Th1 cells. In addition, *Runx1* motifs were enriched within 23 percent of Dnmt3a-dependent DMRs, suggesting that Dnmt3a-mediated methylation may restrict the accessibility of *Runx1* binding sites within the genome. Acquired DNA methylation via Dnmt3a may also directly regulate the *Bcl6* locus in Tfh cells. A previous study by *Lai et al*. demonstrated that DNA methylation inhibited CTCF binding and repression of *Bcl6* at an intronic silencer region within *Bcl6* (Lai et al., 2010). In mouse naïve CD4+ T cells, CTCF is bound to this region (Pham et al., 2019) and our data show that this region acquires *de novo* methylation in Tfh cells. Thus, this mechanism may be conserved between human GC B cells and murine Tfh cells. Collectively, these data strongly implicate Dnmt3a-mediated programing in safeguarding Tfh functionality during the memory recall response by (A) silencing *Runx1*-mediated transcriptional programming and (B) *de novo* methylating an intronic silencer region at the *Bcl6* locus.

In contrast to the *Dnmt3a* knockout model, early treatment with the DNA methyltransferase inhibitor decitabine enhances secondary GC Tfh cell generation and B cell help. Our data suggest that early decitabine treatment during T cell priming partially and temporarily inhibited Dnmt3a-dependent *de novo* methylation at several Tfh-associated genes, including *Cxcr5* and an upstream region of *Bcl6*. Considering that early decitabine treatment induced higher expression of *Bcl6* in Tfh cells, it is plausible that decitabine treatment directly inhibits Dnmt3a-mediated repression of *Bcl6* in Tfh cells, thereby enhancing GC Tfh cell differentiation. Importantly, early decitabine treatment did not inhibit silencing of several loci encoding repressors of Tfh differentiation, including *Prdm1* and *Runx1,* nor the acquisition of *de novo* methylation at the intronic silencer region of *Bcl6* (see previous paragraph). Early decitabine treatment also enhanced the magnitude and function of the secondary Th1 cell response during heterologous rechallenge with Influenza, particularly in the lung. This finding was unexpected considering that decitabine treatment impaired the lineage commitment of memory Th1 cells derived from the spleen. One possible explanation is that decitabine may differentially alter the programing of memory Th1 cell populations in lymphoid and peripheral tissues, but additional research is needed to sort out this question. We cannot rule out that decitabine treatment may also have additional CD4+ T cell-extrinsic effects, which could explain the differences between the decitabine versus knockout models. Dnmt3a-dependent programing regulates gene expression in B cell responses to immunization (Barwick et al., 2018) and CD8 T cell responses to acute and chronic viral infection (Ghoneim et al., 2017, Youngblood et al., 2017, Ladle et al., 2016). We also cannot rule out that decitabine-mediated inhibition of other DNA methyltransferases (Seelan et al., 2018) may contribute in part to the increased GC Tfh phenotype. However, our study demonstrates that (1) decitabine treatment inhibited the acquisition of *de novo* methylation programming at Dnmt3a-dependent regions in virus-specific CD4+ T cells; and (2) that the effects of decitabine treatment were preserved in adoptively-transferred memory T cells following viral challenge of untreated recipient mice. Collectively, these findings strongly support that early decitabine treatment de-represses the Tfh transcriptional program by partial disruption of Dnmt3a-mediated *de novo* methylation, resulting in an intermediate epigenetic program that enhances GC Tfh cell formation and B cell help (Fig. S5F).

Rational vaccine design strategies to improve immunity to viral pathogens have recently focused on increasing the magnitude and duration of the GC response with the objective to enhance selection of neutralizing antibodies and/or increase long-lived plasma cell and memory B cell generation (Crotty, 2019, Yu et al., 2022). Several studies demonstrate that GC Tfh, circulating Tfh, or memory Tfh cell numbers positively correlate with generation of neutralizing antibodies in HIV or Influenza (Yu et al., 2022, Crotty, 2019, Bentebibel et al., 2013, Koutsakos et al., 2018, Locci et al., 2013, Havenar-Daughton et al., 2016, Moody et al., 2016, Cirelli et al., 2019, Lee et al., 2021). In addition, increasing memory Tfh cell numbers accelerates reformation and enhances the magnitude of the GC response following antigen re-encounter (MacLeod et al., 2011, Sircy et al., 2024, Crotty, 2019). Therefore, modulating the abundance and/or functionality of Tfh cells may represent a strategy to enhance the GC response and improve vaccine efficacy (Crotty, 2019). In this study, we demonstrate that early treatment with the methyltransferase inhibitor decitabine during the primary immune response improves Tfh cell abundance and functionality in several ways. First, early decitabine treatment increases primary GC Tfh cell differentiation to several viral pathogens, and in response to influenza infection, this correlates with increased HA-specific GC B cell formation. Second, early decitabine treatment increases memory Tfh cell formation. Third, upon heterologous influenza infection, decitabine treatment during the primary response increases GC Tfh cell recall responses that correlates with increased HA-specific GC B cell responses, without compromising the Th1 cell response in the lung. Collectively, our data demonstrate that DNA methyltransferase inhibition may represent a novel approach to modulate T helper cell differentiation and function to enhance the germinal center response. Altogether, this study identifies key Tfh and Th1 epigenetic programs and novel strategies that could be used to modulate effector and memory T helper cell responses and enhance long-lived immunity to viral pathogens.

## MATERIALS AND METHODS

### Mice

The following mouse strains were purchased from The Jackson Laboratory: C57BL/6J (Strain # 000664), *ERT2-cre* (Strain #008463), *CD4-cre* (Strain #022071), and SMARTA TCR x CD45.1 (Strain #030450). *Dnmt3a^flox^* transgenic mice (Gamper et al., 2009) were bred to generate mice of the following genotypes: (1) *Dnmt3a^flox/flox^* x *ERT2-cre* x SMARTA, (2) *ERT2-cre* x SMARTA, (3) *Dnmt3a^flox/flox^* x *CD4-cre* x SMARTA, (4) *CD4-cre* x SMARTA, and (5) *Dnmt3a^flox/flox^* x SMARTA. All animal experiments were conducted in accordance with University of Utah Institutional Animal Care and Use Committee (IACUC) approved protocols.

### LCMV Armstrong and Influenza Infections

For LCMV experiments, mice were infected with LCMV Armstrong (2x10^5^ pfu) by intraperitoneal injection. For select LCMV experiments that were analyzed at 3 days post infection, mice were infected with LCMV Armstrong (2x10^6^ pfu) by intravenous injection. For all influenza experiments, mice were anesthetized and intranasally infected with either 200 TCID_50_ of the H1N1 PR8 strain or 500 TCID_50_ of the recombinant PR8-GP^61-80^ strain.

### Tissue Processing

Single-cell suspensions of mediastinal lymph nodes or spleens were prepared using 70-μm cell strainers. For spleen processing, red blood cells were lysed by incubation in Ammonium-Chloride-Potassium (ACK) Lysing Buffer (Life Technologies). Single-cell suspensions of lungs were prepared by digestion with 0.25mg/ml Collagenase IV and 15 μg/ml DNase for 1 hour at 37°C, then manually homogenized and red blood cells lysed by incubation in ACK Lysing Buffer and then cells were filtered using 70-μm cell strainers. For all tissues, cell suspensions were resuspended in RPMI 1640 media supplemented with 5% fetal bovine serum (FBS) prior to FACS staining.

### Tamoxifen Treatment and Dnmt3a Knockout Validation

Mice containing the *ERT2-cre* transgene required tamoxifen treatment to induce cre-mediated recombination at the *Dnmt3a* locus. Briefly, mice were intraperitoneally injected with four consecutive doses of tamoxifen (2mg/mouse), followed by a three-day rest period.

Recombination at the *Dnmt3a* locus was validated by qPCR. Briefly, effector SMARTA cells were sorted from splenocytes post infection and DNA was harvested for qPCR analysis. The following primers were used: 5’-TGCAATGACCTCTCCATTGTCAAC-3’, 5’-GGTAGAACTCAAAGAAGAGGCGGC-3’.

Loss of Dnmt3a protein was validated by flow cytometry. Briefly, cells were stained for surface markers prior to fixation and permeabilization. Two step intracellular staining was performed with Rabbit anti-Dnmt3a (D23G1, Cell Signaling Technologies) at 1:100 dilution in perm/wash buffer, followed by AF647-conjugated chicken anti-rabbit IgG (Invitrogen) at 1:500 dilution in perm/wash buffer. A rabbit IgG Isotype control was used as a control.

### Decitabine Treatment

5’-aza-2′-deoxycytidine (decitabine) was purchased from Sigma Aldrich and was dissolved in sterile PBS. For primary LCMV experiments, mice were intraperitoneally injected with decitabine (0.75 mg/kg) 20 hours post primary LCMV infection. For primary influenza experiments, mice were intraperitoneally injected with decitabine (0.375 mg/kg) 20 hours post primary infection.

### Adoptive Transfer of Naïve SMARTA T Cells

Spleens were harvested and processed. For experiments utilizing SMARTA cells without additional transgenes, naïve SMARTA T cells (CD45.1+) were transferred into C57BL/6J mice (CD45.2+). For experiments utilizing *Dnmt3a* knockout cells, naïve SMARTA T cells (CD45.1+) were enriched by negative selection using magnetic beads (Catalog # 19852, STEMCELL Technologies) and then transferred into C57BL/6J, *ERT2-cre,* or *CD4-cre* mice (CD45.2+). The use of *ERT2-cre* or *CD4-cre* containing recipients was specifically for experiments that were analyzed at memory or memory recall timepoints. The number of cells transferred was either 2x10^4^ for analysis at 7 days post infection, 2x10^5^ for memory and memory recall experiments, or 2x10^6^ for analysis at 3 days post infection.

### Adoptive Transfer of Memory SMARTA T Cells

Spleens from LCMV Armstrong infected mice were harvested and processed. The specific times for each experiment are noted in the figure legends. For bulk memory transfer experiments, memory SMARTA T cells were enriched by negative selection using magnetic beads (Catalog # 19852, STEMCELL Technologies), then stained and sorted for CD4+ CD45.1+ memory SMARTA T cells. For experiments that required independent transfer of mTh1 and mTfh populations, splenocytes were first stained with PE-conjugated CD8, CD19, and CD45.2 for negative selection of memory SMARTA T cells using anti-PE MicroBeads (Catalog # 130-048-801, Miltenyi Biotec). Enriched memory SMARTA T cells were then stained and sorted into CXCR5-Ly6C^hi^ mTh1, CXCR5+ Ly6C^hi^ mTfh and CXCR5+ Ly6C^lo^ mTfh populations. For all experiments, 2-10x10^3^ cells were transferred into C57BL/6J mice prior to reinfection with LCMV Armstrong.

### Flow Cytometry and Cell Sorting

All surface staining was done in PBS supplemented with 2% fetal bovine serum (FACS Buffer) for approximately 30 minutes on ice unless specified otherwise. The following antibodies were used for surface staining: CD4 (RM4-5), CD8 (53-6.7), CD44 (IM7), CD45.1 (A20), CD45.2 (104), Ly6C (HK1.4), CXCR5 (L138D7), PD-1 (29F.1A12), ICOS (7E.17G9), CD25 (PC61), CD95 (Jo2), GL7 (GL7), PNA (FL-1071), CD19 (1D3), B220 (RA3-6B2), IgD (11-26c.2a), and CD138 (281-2).

For CXCR5 staining followed by cell sorting, a three-step CXCR5 staining protocol was used as described by *Johnston et al.* (Johnston et al., 2009). For most flow cytometry experiments, CXCR5 was stained using anti-CXCR5 (L138D7) alongside all other surface markers for 30 minutes on ice.

For intracellular staining, cells were fixed and stained using the Foxp3 Permeabilization Fixation kit (eBioscience). The following antibodies were used for intracellular staining: Dnmt3a (D23G1), Bcl6 (K112-91), Tcf1 (S33-966), Tbet (4B10), Blimp1 (5E7), Foxp3 (FJK-16s), Granzyme B (GB12), *Runx1* (RXDMC), Runx2 (D1L7F), Runx3 (R3-5G4), Lef1 (C12A5), Foxo1 (C29H4), and Id2 (ILCID2).

For tetramer staining, cells were stained in RPMI 1640 media supplemented with 10% fetal bovine serum using the following tetramers: I-A^b^:GP^66-77^, I-A^b^:NP^311-325^, or I-A^b^:CLIP^87-101^ (NIH Tetramer Core Facility). Cells were incubated for 1-2 hours at 37°C with 5% CO_2_ prior to surface and intracellular staining steps.

For intracellular cytokine staining, cells were stimulated with GP61–80 peptide in the presence of brefeldin A (BD Biosciences) for 5 hours at 37°C with 5% CO_2_. Following stimulation, cells were surface stained prior to being fixed using the Cytofix Cytoperm kit (BD Biosciences). The following antibodies were used for intracellular staining: IFNγ (XMG1.2), TNFα (MP6-XT22), and IL-2 (JES6-5H4).

For detection of HA-specific B cells, recombinant HA protein from PR8 virus strain (Catalog # IT-003-0010ΔTMp, Immune Technology Corp.) was biotinylated with 80-fold molar excess of NHS-PEG4-Biotin solution from the EZ-Link NHS-PEG4-Biotin kit (Catalog # A39259, ThermoFisher). Excess biotin was removed by buffer exchange of protein into sterile 1X PBS using Zeba Spin Desalting Columns, 7K MWCO (Catalog # 89882, ThermoFisher). Cells were stained on ice for 30min in FACS buffer with 1:100 dilutions of biotin-conjugated-HA and purified rat anti-mouse CD16/CD32 (2.4G2), then stained on ice for 30min in FACS buffer with 1:1000 dilution of APC-conjugated streptavidin.

Flow cytometry experiments were performed on either a FACSCanto or LSRFortessa X-20, and cell sorting experiments were performed on a FACSAria II (BD Biosciences). Data were analyzed using FlowJo software (TreeStar).

### RNA isolation, RNA Sequencing and Analysis

RNA was isolated from sorted CXCR5-Th1 and CXCR5+ Tfh cells using an RNA MiniPrep kit (Zymo Research). RNA-seq libraries were generated using the NEBNext Ultra II Directional RNA Library Kit (New England Biolabs). Prepared libraries were sequenced using an Illumina NovaSeq 6000 system. Sequencing data were aligned to the mm10 reference genome using STAR in two-pass mode to output a BAM file sorted by coordinates. Mapped reads were assigned to annotated genes using featureCounts version 1.6.3, and differentially expressed genes were identified using DESeq2 version 1.30.1 with a 5% false discovery rate.

### DNA Extraction, Whole Genome Methylation Sequencing and Analysis

Genomic DNA was isolated from sorted CXCR5-Th1 and CXCR5+ Tfh cells using a DNeasy Blood and Tissue kit (Qiagen). DNA was fragmented using a S220 Focused Ultrasonicator (Covaris), and sequencing libraries were prepared using the NEBnext Enzymatic Methyl Seq Kit (New England Biolabs). DNA libraries were sequenced using an Illumina NovaSeq 6000 system.

For whole genome bisulfite sequencing, genomic DNA was isolated from sorted effector and memory CXCR5-Ly6C^hi^ Th1 and CXCR5+ Ly6C^lo^ Tfh cells using a DNeasy Blood and Tissue kit (Qiagen). DNA libraries were generated using the NEBNext Ultra II DNA Library Prep kit (New England Biolabs) and bisulfite treatment of the DNA was performed using the EZ DNA Methylation Gold Kit (Zymo Research). DNA libraries were sequenced using an Illumina NovaSeq 6000 system.

For whole genome bisulfite sequencing analysis, sequencing data quality was assessed using FastQC v0.11.4. Adapters were trimmed from the sequencing reads using Trim Galore! v0.4.4 using options (trim_galore -o $OUTDIR --fastqc –paired $FORWARD_READS $REVERSE_READS). Alignment to the mm10 reference genome was performed using Bismark v0.19.0 with options (bismark --multicore 6 --bowtie2 -N 1 $MM10 -1 $FORWARD_READS -2 $REVERSE_READS). Deduplication was performed with deduplicate_bismark (deduplicate_bismark -p -bam $BISMARK_ALIGNED_BAM). Library quality was assessed based on the percentage of reads that aligned to the genome. Library quality was considered sufficient if greater than 50% of reads uniquely aligned to the genome. Enzymatic methyl conversion efficiency was assessed by evaluating the percent of methylation observed in the CHH genome context. Enzymatic methyl conversion was considered sufficient when this value was less than 3%. Genome coverage was assessed using the bedtools genomecov software v2.25.0. Library genome coverage was considered sufficient if 80% of the genome had a depth of at least 10 reads. For each library that met these quality metrics, methylation percentages at individual CpG positions in the reference genome were quantified using the Bismark Methylation Extractor v0.19.0 program with options (bismark_methylation_extractor-p--comprehensive—bedgraph $BISMARK_DEDUPLICATED_BAM). DMRs among the datasets were detected using BSmooth DMR finder. DMRs have at least 10 CpGs and an absolute mean difference of >0.1.

Additional analyses were performed on the whole genome enzymatic methylation sequencing datasets following DMR identification. Heatmaps of the top ranked DMRs were generated using the R package Pheatmap. The specific details on individual ranking metric can be found in the figure legends. Overlapping DMRs were identified using the R package GenomicRanges. Motif analyses were performed using the program HOMER (findMotifsGenome.pl). The following publicly available and processed ChIP sequencing datasets were downloaded from ChIP Atlas (chip-atlas.org): H3K4me1 (SRX1036630) and CTCF (SRX5086069). H3K4me1 (SRX1036630) bigwig files were used to generate heatmaps of signal intensities across DMRs using the Galaxy Server and deepTools (computeMatrix). Human CpG island coordinates at the *Bcl6* locus were collected from *Lai et al.* and were mapped onto the mouse genome (mm10) using LiftOver.

Analysis of specific DMRs was performed using UCSC Genome Browser and included tracks for mammalian conservation and ENCODE identified cis candidate regulatory elements. Grey boxes outlined regions that are differentially methylated. Select DMRs were further analyzed by heatmaps, made in Prism (Graph Pad), of the percent methylation where the column represents individual CpG sites within the region.

### Retroviral Transduction of SMARTA T cells

SMARTA transduction was performed using a modified protocol from *Ye et al.* (Ye et al., 2017). The following MSCV plasmids were ordered from Addgene: *Runx1* - mCherry (Plasmid #80157) and Empty Vector – mCherry (Plasmid #80139). Briefly, HEK293T cells were transduced with a mixture containing either *Runx1* or control plasmids (2ug total DNA), pCL-Eco (1ug, plasmid #12371), TransIT-293 Transfection Reagent (9uL, Mirus Bio), and OptiMEM (200uL, Thermo Fisher Scientific). Transfected HEK293T cells were then incubated for 48 hours at 37°C with 5% CO_2_. One day prior to transduction, SMARTA T cells were preactivated by intravenous injection of GP_61–80_ peptide (200ug/mouse). On the day of transduction, retroviral supernatant was collected and mixed with polybrene (8ug/mL, Sigma Aldrich) and recombinant human IL-2 (10ng/mL, R&D Systems). Preactivated SMARTA T cells enriched by negative selection using magnetic beads (Catalog # 19852, STEMCELL Technologies). SMARTA cells (2x10^6^ cells) were then mixed with retroviral supernatant and spin-transduced for 2 hours at 200xG and 37°C. Transduced SMARTA cells were transferred into C57BL/6J mice prior to LCMV infection.

For intracellular staining of *Runx1* overexpressing effector SMARTA T cells, mCherry+ SMARTA T cells were first surface stained and sorted using a FACSAria II. Cells were subsequently fixed and stained for flow cytometry analysis in the presence of additional congenically marked cells. Additional flow cytometry analyses were performed using bulk splenocytes without fixation to ensure mCherry detection.

### Statistical Analyses

Statistical tests were performed using Prism (Graph Pad).

## ACKNOWLEDGEMENTS

We would like to thank Dr. Matthew Williams (University of Utah) and Dr. Ben Youngblood (St. Jude Children’s Research Hospital) for providing valuable discussion, technical advice, and feedback on the manuscript. We thank the NIH Tetramer Core Facility (Emory University), the University of Utah Flow Cytometry Facility, and the High-Throughput Genomics and Bioinformatic Analysis Shared Resource at Huntsman Cancer Institute at the University of Utah.

This work was supported by NIH grants R01 AI137238 (to J.S.H.) and National Cancer Institute of NIH through awards 5P30CA042014-24 (to University of Utah Flow Cytometry Facility), 1S10RR026802-01 (to University of Utah Flow Cytometry Facility), and P30CA042014 (to University of Utah High-Throughput Genomics and Bioinformatic Analysis).

The authors have no conflicting financial interests.

## ABBREVIATIONS

GC: germinal center
Th1: T helper 1
Tfh: T follicular helper
mTfh: memory Tfh
mTh1: memory
Th1 B6: C57BL/6J
1°: Primary
2°: Secondary
cCRE: cis-candidate regulatory element

---

**Supplemental Figure 1.**
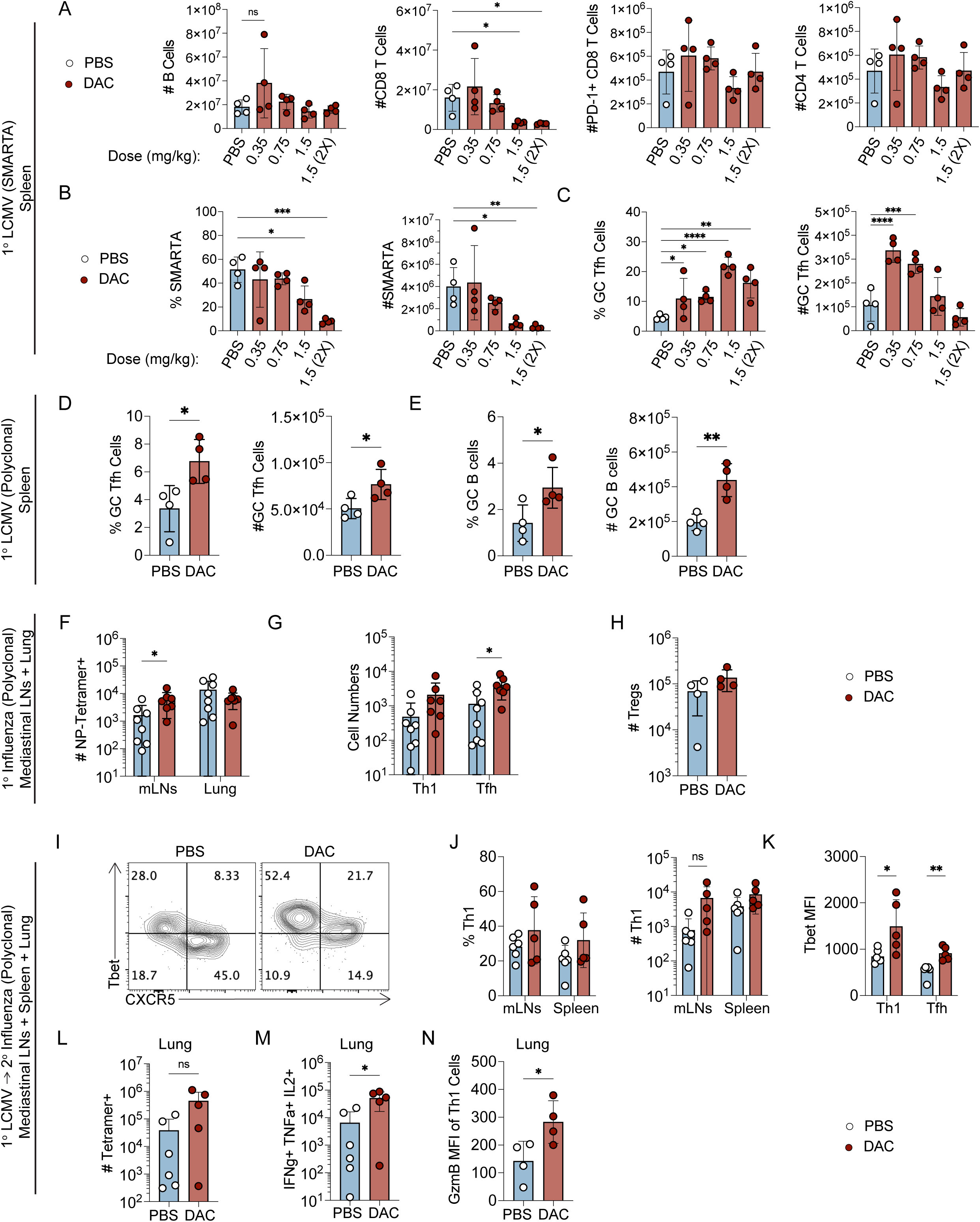
Early decitabine treatment during T cell priming enhances germinal center Tfh differentiation to primary and secondary viral infections. (A-C) Pilot experiment to determine the optimal dose and timing of decitabine treatment. Briefly, naïve SMARTA cells were transferred into B6 mice which were subsequently infected with LCMV. Mice were intraperitoneally injected with either PBS, a single dose of DAC at varying concentrations (0.35mg/kg, 0.75mg/kg, 1.5mg/kg) given 1 day post infection, or two doses of DAC (1.5mg/kg) given on days 1 and 2 post infection. Analysis was performed on the spleen at 7 days p.i. Data are representative of a single experiment. (**A**) Quantification of B cells, CD8 T cells, PD-1+ CD8 T cells and CD4+ T cells. (**B**) Quantification of the percent and numbers of SMARTA T cells and (**C**) CXCR5+ Bcl6^HI^ GC Tfh cells, gated on SMARTA cells. Statistical significance was determined by a one way ANOVA (* p<0.05, **p<0.01, ***p<0.001). (D-E) Briefly, B6 mice which were infected with LCMV. Mice were intraperitoneally injected with either PBS or DAC (0.75mg/kg) at 20 hours post infection. Analysis was performed on the spleen at 7 days p.i. Data are representative of a single experiment. (**D**) Percent and number of PD-1^hi^ GC Tfh cells, gated on CD44^hi^ CD4+ T cells. (**E**) Percent and number of GC B cells, gated on total B cells. Statistical significance was determined by an unpaired t test (* p<0.05, **p<0.01, ***p<0.001). (F-H) B6 mice which were intranasally infected with Influenza PR8. Mice were injected with either PBS or DAC (0.35mg/kg) at 20 hours post infection. Analysis was performed on the mediastinal LNs and lung at 8 days post infection. Data are representative of a two independent experiment. (**F**) Quantification of NP-specific CD4+ T cells in the mediastinal LNs (mLNs) or lung. (**G**) Number of CXCR5-Th1 and CXCR5+ Tfh cells, gated on NP-specific CD4+ T cells. (**H**) Number of Foxp3+ regulatory T cells, gated on CD4+ T cells. Statistical significance was determined by an unpaired t test (* p<0.05, **p<0.01, ***p<0.001). (I-N) B6 mice were infected with LCMV and given a single dose of DAC (0.75mg/kg) 20 hours post infection. At 30+ days p.i., mice were intranasally infected with 500 TCID_50_ of recombinant influenza PR8-GP^61-80^. The following issues were harvested for analysis at 7 days post influenza infection: mediastinal LNs (mLNs), spleen (sp), and lung. Data are representative of two experiments. (**I**) Representative FACS plots of Tbet and CXCR5, gated on GP-specific T cells. (**J**) Quantification of the percent and number of Tbet^hi^ Th1 cells in the mLN and lung. (**K**) Tbet MFI of Tfh and Th1 cells, gated on GP-specific T cells. (**L**) Quantification of NP-specific T cells and (**M**) IFNγ, TNFα, and IL-2 triple producing T cells in the lung, gated on NP-specific T cells. (**N**) Granzyme B (Gzmb) MFI of Th1 cells in the lung, gated on NP-specific T cells.

**Supplemental Figure 2.**
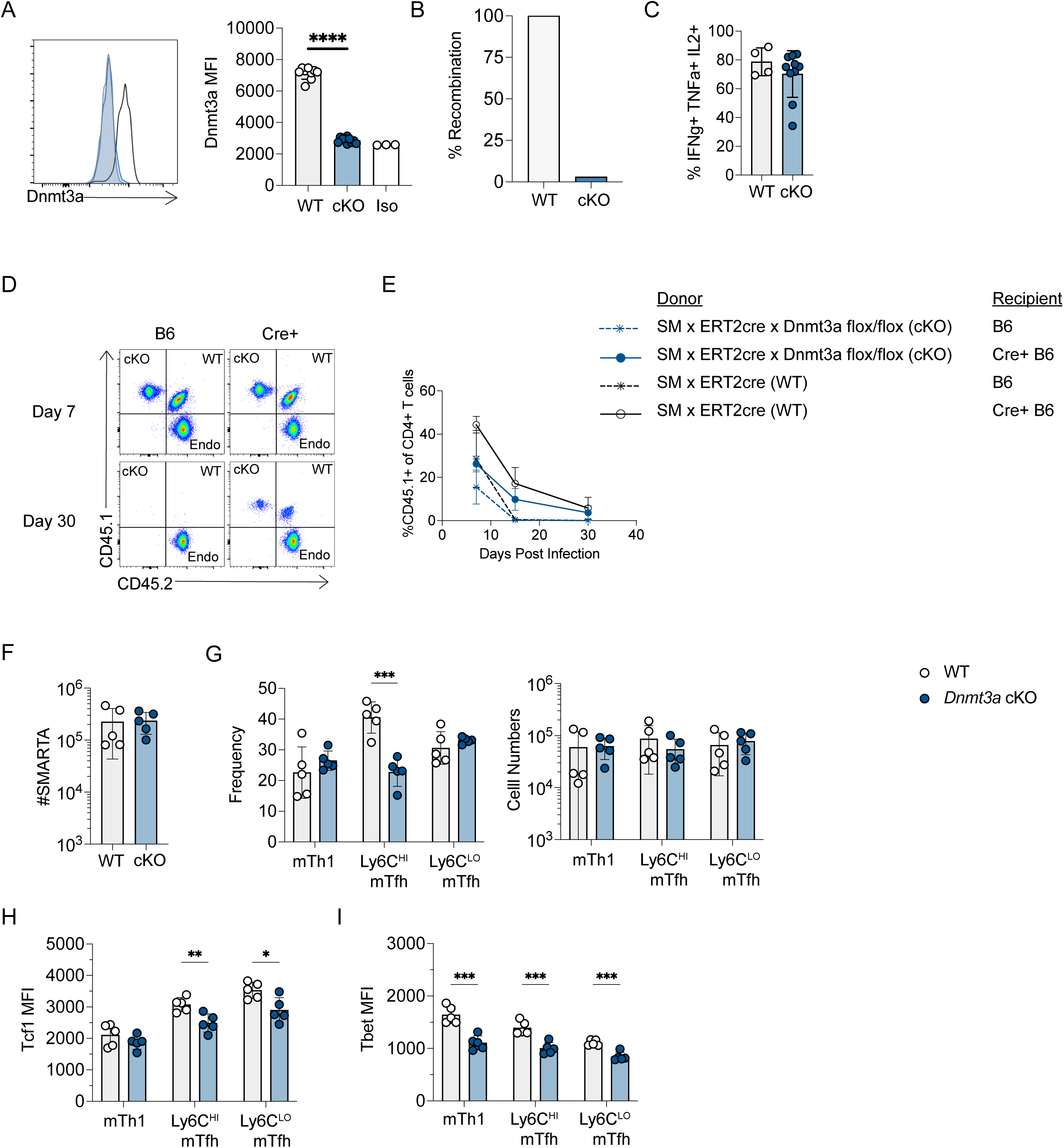
Dnmt3a restricts GC Tfh cell differentiation in a cell-intrinsic manner. (A-C) *ERT2-cre* x SMARTA (WT) or *Dnmt3a^flox/flox^* x *ERT2-cre* x SMARTA (cKO) mice were treated with tamoxifen to induce Dnmt3a deletion. SMARTA cells were transferred into B6 mice and were infected with LCMV. Data was collected from the spleens of mice at 7 days post infection. Data are representative of two independent experiments. (**A**) Representative histogram of Dnmt3a staining and quantification of Dnmt3a MFI gated on SMARTA T cells. (**B**) Quantitative PCR analysis of DNA from either WT or cKO cells to validate recombination of the *Dnmt3a* locus in effector SMARTA cells at 7 days post infection. (**C**) Percent IFNγ, TNFα, and IL-2 triple producers of SMARTA T cells. Statistical significance was determined by an unpaired t test (* p<0.05, **p<0.01, ***p<0.001). (D-E) *ERT2-cre* x SMARTA (WT – CD45.1+ CD45.2+) or *Dnmt3a^flox/flox^* x *ERT2-cre* x SMARTA (cKO – CD45.1+) cells were adoptively transferred into B6 or *ERT2-Cre*+ B6 mice following tamoxifen treatment. Recipients were infected with LCMV, and the CD4+ T cell response was tracked in the blood. Data are representative of three or more experiments. (**D**) Representative FACS plots of CD45.1 and CD45.2 staining for detection of SMARTA T cells, gated on CD4+ T cells. The top row is from 7 days post infection and the bottom is 30 days post infection. (**E**) Quantification of the percent CD45.1+ of CD4+ T cells at 7, 15 and 30 days post infection. (F-I) *ERT2-cre* x SMARTA (WT) or *Dnmt3a^flox/flox^* x *ERT2-cre* x SMARTA (cKO) were adoptively transferred into *ERT2-Cre+* mice and infected with LCMV. Data was collected at 60+ days post infection from the spleens of LCMV infected mice. Data are representative of more than three independent experiments. (**F**) Number of SMARTA cells in the spleen. (**G**) Quantification of the percent and number of CXCR5-Ly6C^HI^ mTh1, CXCR5+ Ly6C^HI^ mTfh, CXCR5+ Ly6C^LO^ mTfh, and CXCR5-Ly6C^LO^ double negative (DN) SMARTA cells. (**H**) Tcf1 MFI and (**I**)Tbet MFI of SMARTA T cell subsets. Statistical significance was determined by an unpaired t test (* p<0.05, **p<0.01, ***p<0.001).

**Supplemental Figure 3.**
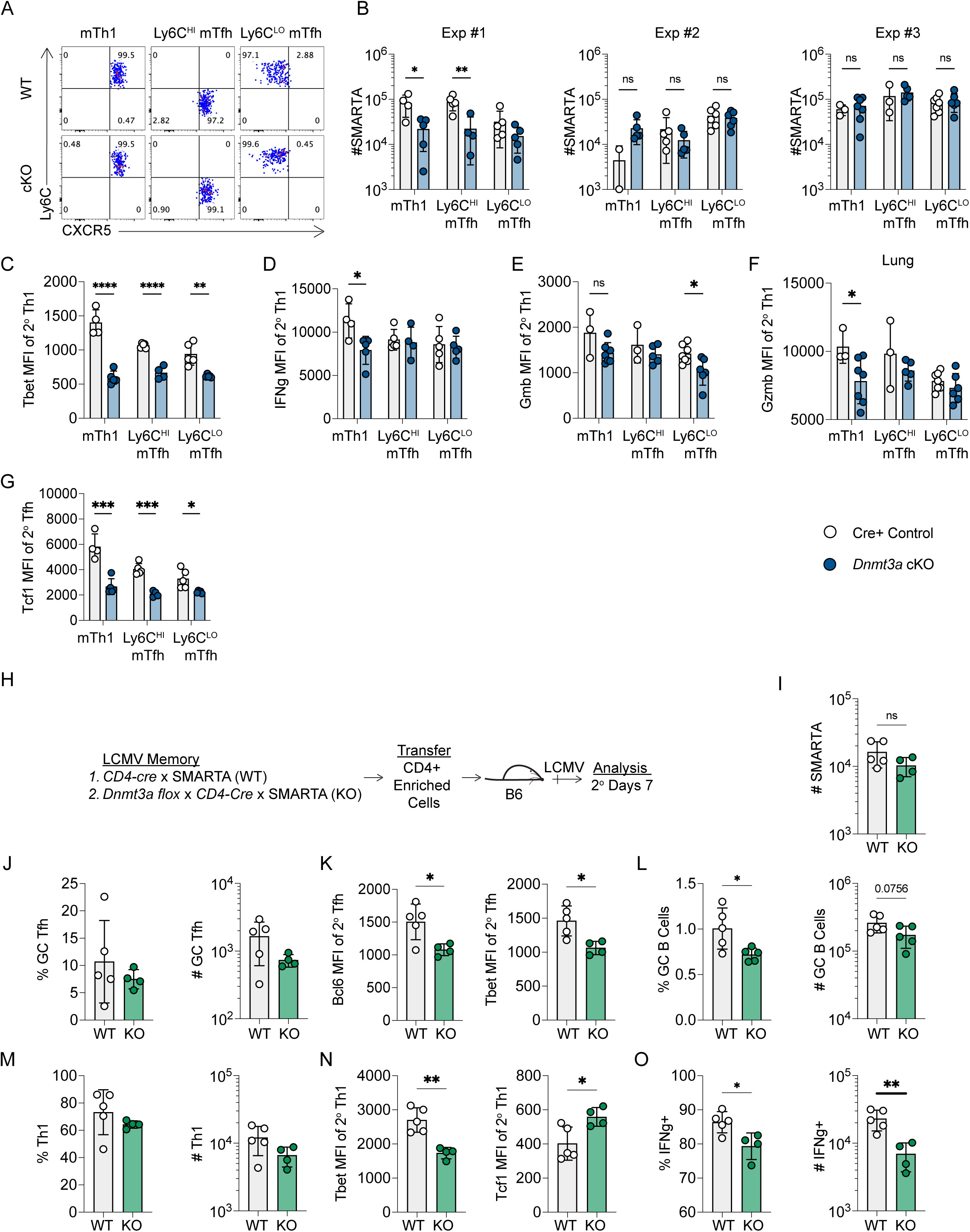
Dnmt3a limits plasticity and preserves functionality of memory Tfh and Th1 cells. (A-G) *ERT2-cre* x SMARTA (WT) or *Dnmt3a^flox/flox^* x *ERT2-cre* x SMARTA (cKO) mice were treated with tamoxifen to induce *Dnmt3a* deletion. SMARTA cells were transferred into B6 mice and were infected with LCMV. At 60+ days post infection, SMARTA cells were sorted from spleens into CXCR5-Ly6C^HI^ mTh1, CXCR5+ Ly6C^HI^ mTfh, and CXCR5+ Ly6C^LO^ mTfh, independently transferred into naïve B6 recipients, and mice were infected with LCMV. The recall response was analyzed in the spleen and lung at 7 days post-secondary infection. Data are representative of three independent experiments. (**A**) Purity analysis of FACS sorted CXCR5-Ly6C^HI^ mTh1, CXCR5+ Ly6C^HI^ mTfh, and CXCR5+ Ly6C^LO^ mTfh cells. (**B**) Quantification of the numbers of SMARTA cells in the spleens across three repeat experiments. Quantifications of (**C**) Tbet MFI of CXCR5-Th1 cells, and (**D**) IFNγ MFI of IFNγ+ cells. Granzyme B MFI of CXCR5-Th1 cells in the (**E**) spleen and (**F**) lung. (**G**) Tcf1 MFIs of CXCR5+ Tfh cells in the spleen. Statistical significance was determined by an unpaired t test (* p<0.05, **p<0.01, ***p<0.001). (H-O) *Dnmt3a^+/+^ x ERT2-cre* (WT) or *Dnmt3a^flox/flox^* x *CD4-cre* (KO) memory SMARTA cells were generated following LCMV infection. At 60+ days post infection, bulk SMARTA memory cells were FACS sorted and transferred into naïve recipients, and mice were infected with LCMV. The recall response was analyzed 7 days post-secondary infection in the spleen. Data are representative of a single experiment. (**H**) Experimental schematic. (**I**) Quantification of the total number of SMARTA cells and (**J**) CXCR5+ Bcl6^HI^ GC Tfh cells, gated on SMARTA cells. (**K**) Quantification of Bcl6 and Tbet MFIs of CXCR5+ Tfh cells. (**L**) Percent and number of GC B cells. (**M**) Percent and number of CXCR5-Th1 cells. (**N**) Tbet MFI and Tcf1 MFI of CXCR5-Th1 cells. (**O**) Percent and number of IFNγ+ of total SMARTA cells. Statistical significance was determined by an unpaired t test (* p<0.05, **p<0.01, ***p<0.001).

**Supplemental Figure 4.**
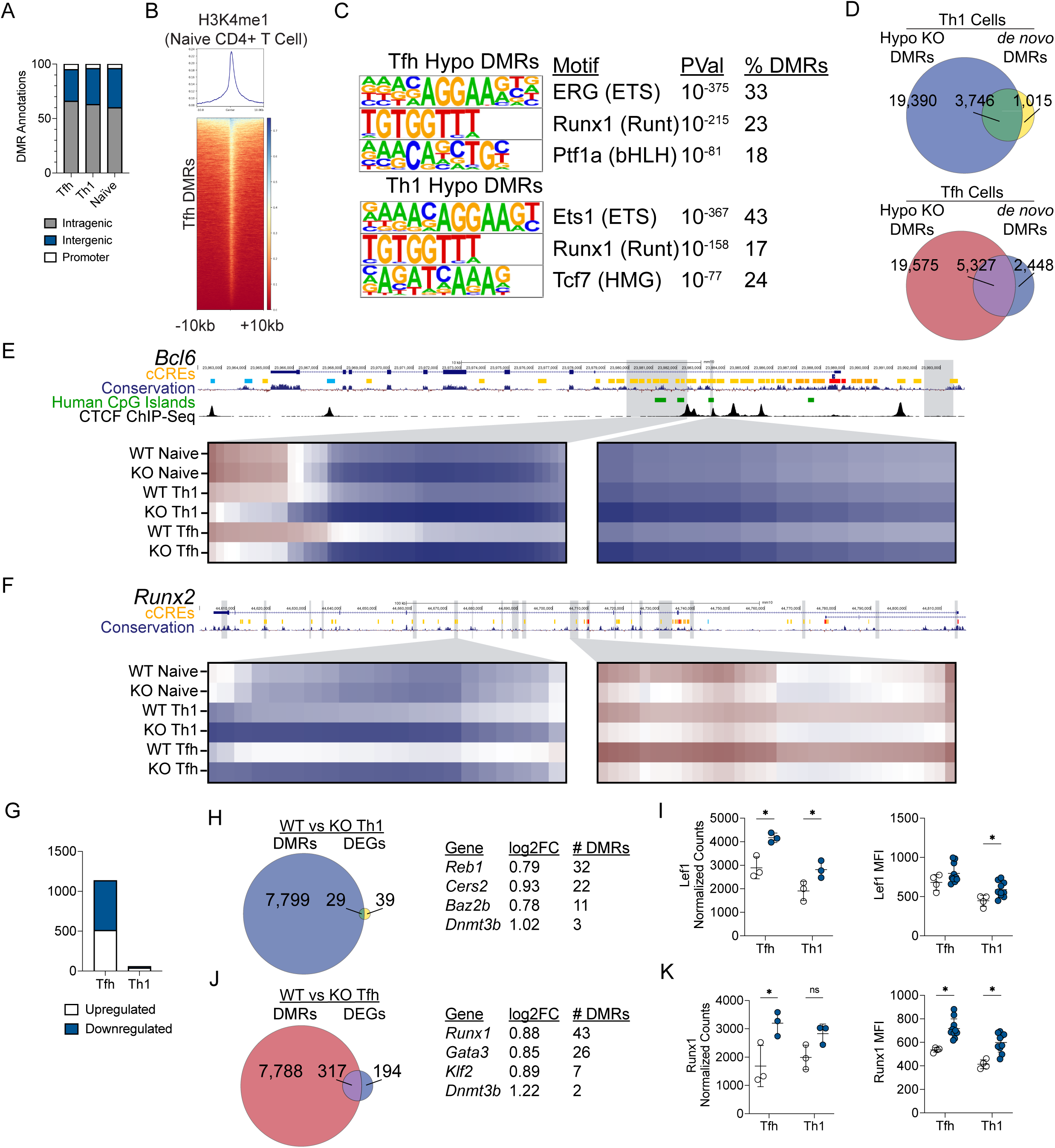
Dnmt3a silences genes associated with alternative T helper lineages in Tfh and Th1 cells. (A-F) Whole genome enzymatic methylation sequencing was performed on *CD4-cre* x SMARTA (WT) and *Dnmt3a^flox/flox^* x *CD4-cre* x SMARTA (KO) Tfh and Th1 cells sorted at 7 days post LCMV infection. WT and KO naïve CD4+ T cells were also analyzed as controls. Differentially methylated regions (DMRs) were called between WT and KO cells of the same lineage. Data are representative of three biological replicates. (**A**) Breakdown of all DMRs by genomic location. (**B**) Enrichment analysis of H3K4me1 ChIP-seq peaks (GSM1694162) from naïve CD4+ T cells. (**C**) Transcription factor motif analysis of Tfh and Th1 DMRs. (**D**) Overlap analysis of Dnmt3a-dependent DMRs and *de novo* DMRs. *De novo* DMRs are regions that are significantly hypermethylated in effector populations relative to naïve CD4+ T cells and are identified in Figure 1. (E-F) UCSC genome browser snapshots of (**E**) *Bcl6,* and (**F**) *Runx2*. We also show tracks for ENCODE identified cis-candidate regulatory elements (cCREs) and mammalian conservation. Grey boxed identify hypomethylated DMRs. In the heatmaps below, select DMRs are shown. Each column represents a CpG site across the DMR. For the *Bcl6* locus only, we also show human CpG islands which were found using LiftOver, as well as a track containing CTCF ChIP seq peaks from Naïve CD4+ T cells (<u>SRX5086069</u>). Significant DMRs were identified using BSmooth DMR finder. DMRs have at least 10 CpGs and an absolute mean difference of >0.1. (G-K) RNA sequencing was performed on *CD4-cre* x SMARTA (WT) and *Dnmt3a^flox/flox^* x *CD4-cre* x SMARTA (KO) Tfh and Th1 cells sorted at 7 days post LCMV infection. Differentially expressed genes (DEGs) were called between WT and cKO cells of the same lineage. Data are representative of three biological replicates. (**G**) Quantification of DEGs, broken down by directionality. (**H**) Overlap analysis of upregulated DEGs that contain hypomethylated DMRs in Th1 cells. (**I**) *Lef1* transcript and protein analysis in effector WT and cKO Tfh/Th1 cells. (**J**) Overlap analysis of upregulated DEGs that contain hypomethylated DMRs in Tfh cells. (**K**) *Runx1* transcript and protein analysis in effector WT and cKO Tfh/Th1 cells. DEGs were identified using DESeq2 version 1.30.1 with a 5% false discovery rate.

**Supplemental Figure 5.**
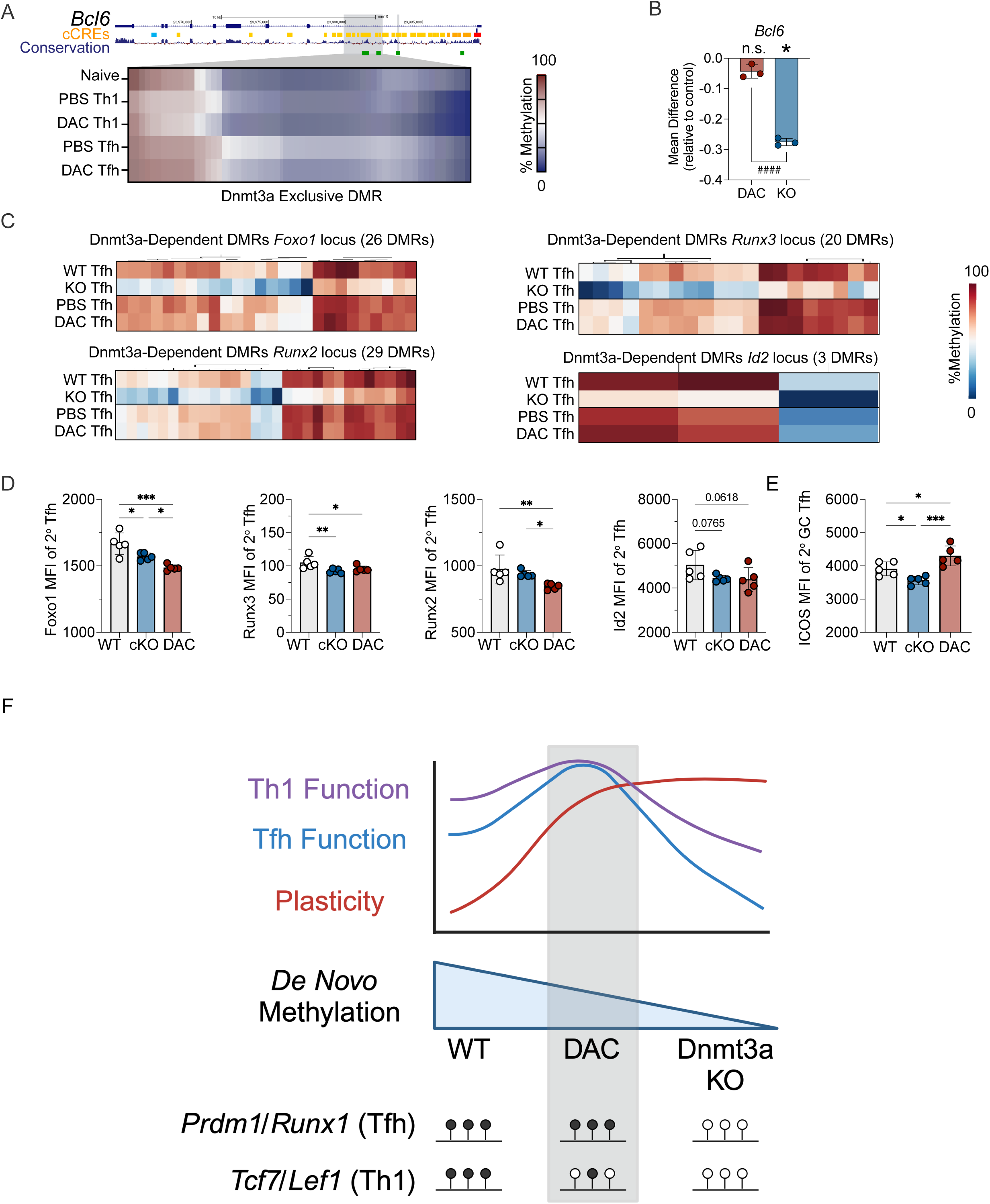
Dnmt3a deficiency, but not early decitabine treatment, impairs silencing of the loci encoding Blimp1 and *Runx1*. (A-C) Additional methylation analysis from Dnmt3a KO or DAC treated samples as shown in Figures 6 and 7. (**A**) UCSC genome browser snapshot of a Dnmt3a-dependent Tfh DMR within the first intron of Bcl6. This select DMR is a Dnmt3a-dependent Tfh DMRs (WT vs KO) and is not a significant DAC-dependent Tfh DMRs (PBS vs DAC). For the locus specific heatmaps, we also show tracks for ENCODE identified cis-candidate regulatory elements (cCREs) and mammalian conservation. Grey boxed identify DMRs, which are highlighted below. Each column represents a CpG site across the region. (**B**) Quantification of the mean difference in methylation for the Dnmt3a-dependent Tfh DMR shown in A. The mean difference for DAC treated Tfh cells and KO Tfh cells are relative to their respective control groups (PBS vs DAC and WT vs KO). (**C**) Heatmaps showing the mean methylation for all Dnmt3a-dependent Tfh DMRs associated with *Foxo1*, *Runx2, Runx3,* and *Id2.* For plots comparing KO and DAC, * symbol represents significant DMR. # symbol represents significant difference between groups as determined by a t test. (D-E) *ERT2-cre* x SMARTA (WT) or *Dnmt3a^flox/flox^* x *ERT2-cre* x SMARTA (cKO) mice were treated with tamoxifen to induce Cre-mediated deletion. Naïve CD4+ T cells were transferred into mice which were subsequently infected with LCMV. Mice were treated with either PBS or DAC (0.75mg/kg) at 20 hours post infection such to generate the following three groups: WT-PBS (WT), cKO-PBS (cKO), and WT-DAC (DAC). At memory timepoint, CD4+ T cells were enriched from the spleen, and transferred into new naïve B6 mice. Mice were infected with LCMV and the SMARTA cell response was analyzed within the spleen at 3- or 7-days post infection. Data are representative of a single experiment. (**D**) Quantification of the MFI of Foxo1, Runx2, Runx3, and Id2. All data are from 2° Tfh SMARTA cells. (**E**) MFI analysis of Icos gated on PD-1^HI^ GC Tfh. Data are from 2° Tfh SMARTA cells. Statistical significance was determined by an unpaired t test (* p<0.05, **p<0.01, ***p<0.001). (**F**) Graphical Abstract.

## REFERENCES

Baessler, A., Fuchs, B., Perkins, B., Richens, A. W., Novis, C. L., HARRISON- Chau, M., Sircy, L. M., Thiede, K. A. C Hale, J. S. 2023. Tet2 deletion in CD4+ T cells disrupts Th1 lineage commitment in memory cells and enhances T follicular helper cell recall responses to viral rechallenge. Proc Natl Acad Sci U S A, 120, e2218324120.

Baessler, A., Novis, C. L., Shen, Z., Perovanovic, J., Wadsworth, M., Thiede, K. A., Sircy, L. M., Harrison-Chau, M., Nguyen, N. X., Varley, K. E., Tantin, D. C Hale, J. S. 2022. Tet2 coordinates with Foxo1 and *Runx1* to balance T follicular helper cell and T helper 1 cell differentiation. Sci Adv, 8, eabm4982.

Barwick, B. G., Scharer, C. D., Martinez, R. J., Price, M. J., Wein, A. N., Haines, R. R., Bally, A. P. R., Kohlmeier, J. E. C Boss, J. M. 2018. B cell activation and plasma cell differentiation are inhibited by *de novo* DNA methylation. Nat Commun, 9, 1900.

Bentebibel, S. E., Lopez, S., Obermoser, G., Schmitt, N., Mueller, C., Harrod, C., Flano, E., Mejias, A., Albrecht, R. A., Blankenship, D., Xu, H., Pascual, V., Banchereau, J., Garcia-Sastre, A., Palucka, A. K., Ramilo, O. C Ueno, H. 2013. Induction of ICOS+CXCR3+CXCR5+ TH cells correlates with antibody responses to influenza vaccination. Sci Transl Med, 5, 176ra32.

Burton, D. R., Ahmed, R., Barouch, D. H., Butera, S. T., Crotty, S., Godzik, A., Kaufmann, D. E., Mcelrath, M. J., Nussenzweig, M. C., Pulendran, B., Scanlan, C. N., Schief, W. R., Silvestri, G., Streeck, H., Walker, B. D., Walker, L. M., Ward, A. B., Wilson, I. A. C Wyatt, R. 2012. A Blueprint for HIV Vaccine Discovery. Cell Host Microbe, 12, 396–407.

Choi, J. C Crotty, S. 2021. Bcl6-Mediated Transcriptional Regulation of Follicular Helper T cells (T(FH)). Trends Immunol, 42, 336–349.

Choi, J., Diao, H., Faliti, C. E., Truong, J., Rossi, M., Bélanger, S., Yu, B., Goldrath, A. W., Pipkin, M. E. C Crotty, S. 2020. Bcl-6 is the nexus transcription factor of T follicular helper cells via repressor-of-repressor circuits. Nat Immunol, 21, 777–789.

Choi, Y. S., Gullicksrud, J. A., Xing, S., Zeng, Z., Shan, Q., Li, F., Love, P. E., Peng, W., Xue, H. H. C Crotty, S. 2015. LEF-1 and TCF-1 orchestrate T(FH) differentiation by regulating differentiation circuits upstream of the transcriptional repressor Bcl6. Nat Immunol, 16, 980–90.

Choi, Y. S., Yang, J. A., Yusuf, I., Johnston, R. J., Greenbaum, J., Peters, B. C Crotty, S. 2013. Bcl6 expressing follicular helper CD4 T cells are fate committed early and have the capacity to form memory. J Immunol, 190, 4014–26.

Christman, J. K. 2002. 5-Azacytidine and 5-aza-2’-deoxycytidine as inhibitors of DNA methylation: mechanistic studies and their implications for cancer therapy. Oncogene, 21, 5483–95.

Cirelli, K. M., Carnathan, D. G., Nogal, B., Martin, J. T., Rodriguez, O. L., Upadhyay, A. A., Enemuo, C. A., Gebru, E. H., Choe, Y., Viviano, F., Nakao, C., Pauthner, M. G., Reiss, S., Cottrell, C. A., Smith, M. L., Bastidas, R., Gibson, W., Wolabaugh, A. N., Melo, M. B., Cossette, B., Kumar, V., Patel, N. B., Tokatlian, T., Menis, S., Kulp, D. W., Burton, D. R., Murrell, B., Schief, W. R., Bosinger, S. E., Ward, A. B., Watson, C. T., Silvestri, G., Irvine, D. J. C Crotty, S. 2019. Slow Delivery Immunization Enhances HIV Neutralizing Antibody and Germinal Center Responses via Modulation of Immunodominance. Cell, 177, 1153–1171.e28.

Crotty, S. 2019. T Follicular Helper Cell Biology: A Decade of Discovery and Diseases. Immunity, 50, 1132–1148.

Derissen, E. J., Beijnen, J. H. C Schellens, J. H. 2013. Concise drug review: azacitidine and decitabine. Oncologist, 18, 619–24.

Feng, H., Zhao, Z., Zhao, X., Bai, X., Fu, W., Zheng, L., Kang, B., Wang, X., Zhang, Z. C Dong, C. 2024. A novel memory-like Tfh cell subset is precursor to effector Tfh cells in recall immune responses. J Exp Med, 221.

Gamper, C. J., Agoston, A. T., Nelson, W. G. C Powell, J. D. 2009. Identification of DNA methyltransferase 3a as a T cell receptor-induced regulator of Th1 and Th2 differentiation. J Immunol, 183, 2267–76.

Ghoneim, H. E., Fan, Y., Moustaki, A., Abdelsamed, H. A., Dash, P., Dogra, P., Carter, R., Awad, W., Neale, G., Thomas, P. G. C Youngblood, B. 2017. De Novo Epigenetic Programs Inhibit PD-1 Blockade-Mediated T Cell Rejuvenation. Cell, 170, 142–157.e19.

Greenberg, M. V. C. C BOURC’his, D. 2019. The diverse roles of DNA methylation in mammalian development and disease. Nat Rev Mol Cell Biol, 20, 590–607.

Hale, J. S., Youngblood, B., Latner, D. R., Mohammed, A. U., Ye, L., Akondy, R. S., Wu, T., Iyer, S. S. C Ahmed, R. 2013. Distinct memory CD4+ T cells with commitment to T follicular helper- and T helper 1-cell lineages are generated after acute viral infection. Immunity, 38, 805–17.

Hatzi, K., Nance, J. P., Kroenke, M. A., Bothwell, M., Haddad, E. K., Melnick, A. C Crotty, S. 2015. BCL6 orchestrates Tfh cell differentiation via multiple distinct mechanisms. J Exp Med, 212, 539–53.

Havenar-Daughton, C., Carnathan, D. G., Torrents De la PEña, A., Pauthner, M., Briney, B., Reiss, S. M., Wood, J. S., Kaushik, K., VAN Gils, M. J., Rosales, S. L., VAN DER Woude, P., Locci, M., Le, K. M., DE Taeye, S. W., Sok, D., Mohammed, A. U. R., Huang, J., Gumber, S., Garcia, A., Kasturi, S. P., Pulendran, B., Moore, J. P., Ahmed, R., Seumois, G., Burton, D. R., Sanders, R. W., Silvestri, G. C Crotty, S. 2016. Direct Probing of Germinal Center Responses Reveals Immunological Features and Bottlenecks for Neutralizing Antibody Responses to HIV Env Trimer. Cell Rep, 17, 2195–2209.

Ichiyama, K., Chen, T., Wang, X., Yan, X., Kim, B. S., Tanaka, S., Ndiaye-Lobry, D., Deng, Y., Zou, Y., Zheng, P., Tian, Q., Aifantis, I., Wei, L. C Dong, C. 2015. The methylcytosine dioxygenase Tet2 promotes DNA demethylation and activation of cytokine gene expression in T cells. Immunity, 42, 613–26.

Johnston, R. J., Poholek, A. C., Ditoro, D., Yusuf, I., Eto, D., Barnett, B., Dent, A. L., Craft, J. C Crotty, S. 2009. Bcl6 and Blimp-1 are reciprocal and antagonistic regulators of T follicular helper cell differentiation. Science, 325, 1006–10.

Koutsakos, M., Wheatley, A. K., Loh, L., Clemens, E. B., Sant, S., Nüssing, S., Fox, A., Chung, A. W., Laurie, K. L., Hurt, A. C., Rockman, S., Lappas, M., Loudovaris, T., Mannering, S. I., Westall, G. P., Elliot, M., Tangye, S. G., Wakim, L. M., Kent, S. J., Nguyen, T. H. O. C Kedzierska, K. 2018. Circulating T(FH) cells, serological memory, and tissue compartmentalization shape human influenza-specific B cell immunity. Sci Transl Med, 10.

Krammer, F. C Palese, P. 2015. Advances in the development of influenza virus vaccines. Nat Rev Drug Discov, 14, 167–82.

Künzli, M. C Masopust, D. 2023. CD4(+) T cell memory. Nat Immunol, 24, 903–914.

Künzli, M., Schreiner, D., Pereboom, T. C., Swarnalekha, N., Litzler, L. C., Lötscher, J., Ertuna, Y. I., Roux, J., Geier, F., Jakob, R. P., Maier, T., Hess, C., Taylor, J. J. C King, C. G. 2020. Long-lived T follicular helper cells retain plasticity and help sustain humoral immunity. Sci Immunol, 5.

Ladle, B. H., Li, K. P., Phillips, M. J., Pucsek, A. B., Haile, A., Powell, J. D., Jaffee, E. M., Hildeman, D. A. C Gamper, C. J. 2016. De novo DNA methylation by DNA methyltransferase 3a controls early effector CD8+ T-cell fate decisions following activation. Proc Natl Acad Sci U S A, 113, 10631–6.

Lai, A. Y., Fatemi, M., Dhasarathy, A., Malone, C., Sobol, S. E., Geigerman, C., Jaye, D. L., Mav, D., Shah, R., Li, L. C Wade, P. A. 2010. DNA methylation prevents CTCF-mediated silencing of the oncogene BCL6 in B cell lymphomas. J Exp Med, 207, 1939–50.

Lai, C. Y., Marcel, N., Yaldiko, A. W., Delpoux, A. C Hedrick, S. M. 2022. A Bcl6 Intronic Element Regulates T Follicular Helper Cell Differentiation. J Immunol, 209, 1118–1127.

Lee, J. H., Hu, J. K., Georgeson, E., Nakao, C., Groschel, B., Dileepan, T., Jenkins, M. K., Seumois, G., Vijayanand, P., Schief, W. R. C Crotty, S. 2021. Modulating the quantity of HIV Env-specific CD4 T cell help promotes rare B cell responses in germinal centers. J Exp Med, 218.

Liu, X., Yan, X., Zhong, B., Nurieva, R. I., Wang, A., Wang, X., Martin-Orozco, N., Wang, Y., Chang, S. H., Esplugues, E., Flavell, R. A., Tian, Q. C Dong, C. 2012. Bcl6 expression specifies the T follicular helper cell program in vivo. J Exp Med, 209, 1841–52, s1-24.

Locci, M., Havenar-Daughton, C., Landais, E., Wu, J., Kroenke, M. A., Arlehamn, C. L., Su, L. F., Cubas, R., Davis, M. M., Sette, A., Haddad, E. K., Poignard, P. C Crotty, S. 2013. Human circulating PD-1+CXCR3-CXCR5+ memory Tfh cells are highly functional and correlate with broadly neutralizing HIV antibody responses. Immunity, 39, 758–69.

Lüthje, K., Kallies, A., Shimohakamada, Y., Belz, G. T., Light, A., Tarlinton, D. M. C Nutt, S. L. 2012. The development and fate of follicular helper T cells defined by an IL-21 reporter mouse. Nat Immunol, 13, 491–8.

Macleod, M. K., David, A., Mckee, A. S., Crawford, F., Kappler, J. W. C Marrack, P. 2011. Memory CD4 T cells that express CXCR5 provide accelerated help to B cells. J Immunol, 186, 2889–96.

Marshall, H. D., Chandele, A., Jung, Y. W., Meng, H., Poholek, A. C., Parish, I. A., Rutishauser, R., Cui, W., Kleinstein, S. H., Craft, J. C Kaech, S. M. 2011. Differential expression of Ly6C and T-bet distinguish effector and memory Th1 CD4(+) cell properties during viral infection. Immunity, 35, 633–46.

Moody, M. A., Pedroza-Pacheco, I., Vandergrift, N. A., Chui, C., Lloyd, K. E., Parks, R., Soderberg, K. A., Ogbe, A. T., Cohen, M. S., Liao, H. X., Gao, F., Mcmichael, A. J., Montefiori, D. C., Verkoczy, L., Kelsoe, G., Huang, J., Shea, P. R., Connors, M., Borrow, P. C Haynes, B. F. 2016. Immune perturbations in HIV-1-infected individuals who make broadly neutralizing antibodies. Sci Immunol, 1, aag0851.

Morita, R., Schmitt, N., Bentebibel, S. E., Ranganathan, R., Bourdery, L., Zurawski, G., Foucat, E., Dullaers, M., Oh, S., Sabzghabaei, N., Lavecchio, E. M., Punaro, M., Pascual, V., Banchereau, J. C Ueno, H. 2011. Human blood CXCR5(+)CD4(+) T cells are counterparts of T follicular cells and contain specific subsets that differentially support antibody secretion. Immunity, 34, 108–21.

Nurieva, R. I., Chung, Y., Martinez, G. J., Yang, X. O., Tanaka, S., Matskevitch, T. D., Wang, Y. H. C Dong, C. 2009. Bcl6 mediates the development of T follicular helper cells. Science, 325, 1001–5.

Oxenius, A., Bachmann, M. F., Zinkernagel, R. M. C Hengartner, H. 1998. Virus-specific MHC-class II-restricted TCR-transgenic mice: effects on humoral and cellular immune responses after viral infection. Eur J Immunol, 28, 390–400.

Pepper, M., PAGán, A. J., IGYáRTó, B. Z., Taylor, J. J. C Jenkins, M. K. 2011. Opposing signals from the Bcl6 transcription factor and the interleukin-2 receptor generate T helper 1 central and effector memory cells. Immunity, 35, 583–95.

Pham, D., Moseley, C. E., Gao, M., Savic, D., Winstead, C. J., Sun, M., Kee, B. L., Myers, R. M., Weaver, C. T. C Hatton, R. D. 2019. Batf Pioneers the Reorganization of Chromatin in Developing Effector T Cells via Ets1-Dependent Recruitment of Ctcf. Cell Rep, 29, 1203–1220.e7.

Pham, D., Yu, Q., Walline, C. C., Muthukrishnan, R., Blum, J. S. C Kaplan, M. H. 2013. Opposing roles of STAT4 and Dnmt3a in Th1 gene regulation. J Immunol, 191, 902–11.

Placek, K., Hu, G., Cui, K., Zhang, D., Ding, Y., Lee, J. E., Jang, Y., Wang, C., Konkel, J. E., Song, J., Liu, C., Ge, K., Chen, W. C Zhao, K. 2017. MLL4 prepares the enhancer landscape for Foxp3 induction via chromatin looping. Nat Immunol, 18, 1035–1045.

Seelan, R. S., Mukhopadhyay, P., Pisano, M. M. C Greene, R. M. 2018. Effects of 5-Aza-2’-deoxycytidine (decitabine) on gene expression. Drug Metab Rev, 50, 193–207.

Shao, P., Li, F., Wang, J., Chen, X., Liu, C. C Xue, H. H. 2019. Cutting Edge: Tcf1 Instructs T Follicular Helper Cell Differentiation by Repressing Blimp1 in Response to Acute Viral Infection. J Immunol, 203, 801–806.

Sircy, L. M., Ramstead, A. G., Gibbs, L. C., Joshi, H., Baessler, A., Mena, I., García-Sastre, A., Emerson, L. L., Fairfax, K. C., Williams, M. A. C Hale, J. S. 2024. Generation of antigen-specific memory CD4 T cells by heterologous immunization enhances the magnitude of the germinal center response upon influenza infection. PLoS Pathog, 20, e1011639.

Stone, E. L., Pepper, M., Katayama, C. D., Kerdiles, Y. M., Lai, C. Y., Emslie, E., Lin, Y. C., Yang, E., Goldrath, A. W., Li, M. O., Cantrell, D. A. C Hedrick, S. M. 2015. ICOS coreceptor signaling inactivates the transcription factor FOXO1 to promote Tfh cell differentiation. Immunity, 42, 239–251.

Thomas, R. M., Gamper, C. J., Ladle, B. H., Powell, J. D. C Wells, A. D. 2012. De novo DNA methylation is required to restrict T helper lineage plasticity. J Biol Chem, 287, 22900–9.

Tsagaratou, A., ÄIJö, T., Lio, C. W., Yue, X., Huang, Y., Jacobsen, S. E., LäHDESMäki, H. C Rao, A. 2014. Dissecting the dynamic changes of 5-hydroxymethylcytosine in T-cell development and differentiation. Proc Natl Acad Sci U S A, 111, E3306–15.

Weber, J. P., Fuhrmann, F. C Hutloff, A. 2012. T-follicular helper cells survive as long-term memory cells. Eur J Immunol, 42, 1981–8.

Wong, W. F., Kohu, K., Nakamura, A., Ebina, M., Kikuchi, T., Tazawa, R., Tanaka, K., Kon, S., Funaki, T., Sugahara-Tobinai, A., Looi, C. Y., Endo, S., Funayama, R., Kurokawa, M., Habu, S., Ishii, N., Fukumoto, M., Nakata, K., Takai, T. C Satake, M. 2012. Runx1 deficiency in CD4+ T cells causes fatal autoimmune inflammatory lung disease due to spontaneous hyperactivation of cells. J Immunol, 188, 5408–20.

Wu, T., Shin, H. M., Moseman, E. A., Ji, Y., Huang, B., Harly, C., Sen, J. M., Berg, L. J., Gattinoni, L., Mcgavern, D. B. C Schwartzberg, P. L. 2015. TCF1 Is Required for the T Follicular Helper Cell Response to Viral Infection. Cell Rep, 12, 2099–110.

Xu, L., Cao, Y., Xie, Z., Huang, Q., Bai, Q., Yang, X., He, R., Hao, Y., Wang, H., Zhao, T., Fan, Z., Qin, A., Ye, J., Zhou, X., Ye, L. C Wu, Y. 2015. The transcription factor TCF-1 initiates the differentiation of T(FH) cells during acute viral infection. Nat Immunol, 16, 991–9.

Ye, L., Lee, J., Xu, L., Mohammed, A. U., Li, W., Hale, J. S., Tan, W. G., Wu, T., Davis, C. W., Ahmed, R. C Araki, K. 2017. mTOR Promotes Antiviral Humoral Immunity by Differentially Regulating CD4 Helper T Cell and B Cell Responses. J Virol, 91.

Youngblood, B., Hale, J. S., Kissick, H. T., Ahn, E., Xu, X., Wieland, A., Araki, K., West, E. E., Ghoneim, H. E., Fan, Y., Dogra, P., Davis, C. W., Konieczny, B. T., Antia, R., Cheng, X. C Ahmed, R. 2017. Effector CD8 T cells dedifferentiate into long-lived memory cells. Nature, 552, 404–409.

Yu, D., Rao, S., Tsai, L. M., Lee, S. K., He, Y., Sutcliffe, E. L., Srivastava, M., Linterman, M., Zheng, L., Simpson, N., Ellyard, J. I., Parish, I. A., Ma, C. S., Li, Q. J., Parish, C. R., Mackay, C. R. C Vinuesa, C. G. 2009. The transcriptional repressor Bcl-6 directs T follicular helper cell lineage commitment. Immunity, 31, 457–68.

Yu, D., Walker, L. S. K., Liu, Z., Linterman, M. A. C Li, Z. 2022. Targeting T(FH) cells in human diseases and vaccination: rationale and practice. Nat Immunol, 23, 1157–1168.

Yu, Q., Zhou, B., Zhang, Y., Nguyen, E. T., Du, J., Glosson, N. L. C Kaplan, M. H. 2012. DNA methyltransferase 3a limits the expression of interleukin-13 in T helper 2 cells and allergic airway inflammation. Proc Natl Acad Sci U S A, 109, 541–6.

Yue, X., Lio, C. J., Samaniego-Castruita, D., Li, X. C Rao, A. 2019. Loss of TET2 and TET3 in regulatory T cells unleashes effector function. Nat Commun, 10, 2011.

Yue, X., Samaniego-Castruita, D., González-Avalos, E., Li, X., Barwick, B. G. C Rao, A. 2021. Whole-genome analysis of TET dioxygenase function in regulatory T cells. EMBO Rep, 22, e52716.

Yue, X., Trifari, S., ÄIJö, T., Tsagaratou, A., Pastor, W. A., Zepeda-Martínez, J. A., Lio, C. W., Li, X., Huang, Y., Vijayanand, P., LäHDESMäki, H. C Rao, A. 2016. Control of Foxp3 stability through modulation of TET activity. J Exp Med, 213, 377–97.

Zhong, Y., Walker, S. K., Pritykin, Y., Leslie, C. S., Rudensky, A. Y. C Van Der Veeken, J. 2022. Hierarchical regulation of the resting and activated T cell epigenome by major transcription factor families. Nat Immunol, 23, 122–134.

Zhu, J., Yamane, H. C Paul, W. E. 2010. Differentiation of effector CD4 T cell populations (*). Annu Rev Immunol, 28, 445–89.

